# Two distinct HIRA-dependent pathways handle H3.3 *de novo* deposition and recycling during transcription

**DOI:** 10.1101/2019.12.18.880716

**Authors:** Júlia Torné, Dominique Ray-Gallet, Ekaterina Boyarchuk, Mickaël Garnier, Antoine Coulon, Guillermo A. Orsi, Geneviève Almouzni

## Abstract

The packaging of DNA into nucleosomes represents a challenge for transcription. Nucleosome disruption and histone eviction enables RNA Polymerase II progression through DNA, a process that compromises chromatin integrity and the maintenance of epigenetic information. Here, we used the imaging SNAP-tag system to distinguish new and old histones and monitor chromatin re-assembly coupled to transcription in cells. First, we uncovered a loss of both old variants H3.1 and H3.3 that depends on transcriptional activity, with a major effect on H3.3. Focusing on transcriptionally active domains, we revealed a local enrichment in H3.3 with dynamics involving both new H3.3 incorporation and old H3.3 retention. Mechanistically, we demonstrate that the HIRA chaperone is critical to handle both new and old H3.3, and showed that this implicates different pathways. The de novo H3.3 deposition depends strictly on HIRA trimerization as well as its partner UBN1 while ASF1 interaction with HIRA can be bypassed. In contrast, the recycling of H3.3 requires HIRA but proceeds independently of UBN1 or HIRA trimerization and shows an absolute dependency on ASF1-HIRA interaction. Therefore, we propose a model where HIRA can coordinate these distinct pathways for old H3.3 recycling and new H3.3 deposition during transcription to fine-tune chromatin states.

## INTRODUCTION

Transcription is intimately linked to chromatin organization. The basic unit of chromatin, the nucleosome, comprises an octamer of four histones – a tetramer (H3-H4)_2_, flanked by two dimers of H2A-H2B – around which 146 bp of DNA is wrapped, plus additional linker DNA (Luger 1997). Nucleosomes have been reported to be a barrier to transcription initiation and to interfere with RNA Polymerase II (RNAPII) progression (Knezetic and Luse 1986; Lorch *et al*. 1987). Thus, nucleosomes are readily disrupted and mobilized during transcription *in vivo* to allow mRNA synthesis (Janicki *et al*. 2004; Thiriet and Hayes 2005; Petesch and Lis 2008). Furthermore, nucleosomes are not monotonous entities; they contain distinct histone variants with the addition of post-translational modifications (PTMs). These chromatin features define different chromosomal landmarks and influence cell fate (Talbert and Henikoff 2010; Yadav *et al*. 2018). In this context, it is particularly crucial to understand how the stable maintenance of chromatin profiles reconciles with the fact that transcription itself is a disruptive process for chromatin.

Mechanistically, torsional stress of DNA caused by the transcription bubble favours nucleosome disassembly downstream from RNAPII (Teves and Henikoff 2014). In addition, ATP-dependent chromatin remodellers and histone chaperones can also displace or disassemble nucleosomes ahead of RNAPII (Venkatesh and Workman 2015). As a result of these disruptive dynamics, active genes are usually characterized by low nucleosome occupancy (Lee *et al*. 2004; Baldi *el al.* 2018) and high rates of histone turnover (Dion *et al*. 2007; Deal *et al*. 2010; Kraushaar *et al*. 2013; Deaton *et al*. 2016). Yet, *in vitro* studies have revealed that RNA Polymerase can bypass nucleosomes at low transcription rates (Kireeva *et al*. 2002; 2005; Bondarenko *et al*. 2006; Kulaeva *et al*. 2009; 2010; Kujirai *et al*. 2018). Furthermore, recent advances in cryo-EM technology have enabled the snapshot structure of the RNAPII-nucleosome complex; providing a framework to envisage nucleosome mobilization and recycling in the wake of transcription (Farnung *et al*. 2018; Kujirai *et al*. 2018). At this point in time, a major issue is thus to which extent chromatin states are actually altered by transcription (Kujirai and Kurumizaka, 2019). However, it is clear that the epigenomic landscape at actively transcribing domains is constantly challenged, compromising chromatin integrity and the maintenance of epigenomic information.

Counteracting these disruptive events, mechanisms that coordinate chromatin assembly coupled to transcription have been reported (Schwabish and Struhl 2004; Schwartz and Ahmad 2005). These possibly involve either *de novo* deposition of new histones, recycling of pre-existing (old) ones, or a combination of both. To date, significant progress has been made to understand mechanisms involved in *de novo* deposition for newly synthesized histone variants. This is best exemplified by the transcription-dependent replacement of the replicative histone H3.1 with the histone variant H3.3 (Janicki *et al*. 2004; Schwartz and Ahmad 2005). New deposition of the replicative histone H3.1 –which constitutes the bulk of histone H3 in proliferative cells – involves a DNA synthesis coupled (DSC) pathway (Ahmad and Henikoff 2002; Tagami *et al*. 2004) that depends on the Chromatin Assembly Factor 1 (CAF-1) complex (Tagami et al. 2004). In contrast, new deposition of H3.3 proceeds in a DNA synthesis independent (DSI) manner (Ahmad and Henikoff 2002; Tagami *et al*. 2004). Excepted for its accumulation in heterochromatic regions, which involves the death domain-associated protein *α*-thalassaemia/mental retardation syndrome X-linked (DAXX-ATRX) complex (Drane *et al*. 2010; Wong *et al*. 2010; Lewis *et al*. 2010; Goldberg *et al*. 2010), most new H3.3 deposition depends on the Histone regulator A (HIRA) chaperone pathway (Tagami *et al*. 2004; Ray-Gallet *et al*. 2011).

At active genes, H3.3 is specifically enriched in a HIRA-dependent manner (Goldberg *et al*. 2010; Ray-Gallet *et al*. 2011). Importantly, HIRA is part of a complex comprising three distinct polypeptides (Tagami *et al*. 2004): HIRA itself as a scaffold protein, Calcineurin binding protein 1 (CABIN1) (Rai *et al*. 2011) and Ubinuclein 1 (UBN1) (Banumathy *et al*. 2009). UBN1 is essential for *de novo* histone deposition through a direct interaction with H3.3-H4 dimers that enables their transfer onto DNA (Ray-Gallet *et al*. 2011; Orsi *et al*. 2013; Ricketts *et al*. 2015; 2019). The HIRA subunit homotrimerizes and associates with two CABIN1 subunits (Ray-Gallet *et al*. 2018). Notably, HIRA trimerization is necessary for *de novo* deposition of H3.3 (Ray-Gallet *et al*. 2018). In addition to these core partners, HIRA can also interact with the Anti-silencing Function 1 (ASF1) histone chaperone (Tang *et al*. 2006; Green *et al*. 2010). ASF1b or ASF1a escort H3.1-H4 and H3.3-H4 dimers and can hand these dimers off respectively to either CAF-1 or HIRA complexes (Tyler *et al*. 1999; Mello *et al*. 2002; Daganzo *et al*. 2003; English *et al*. 2006; Cook *et al*. 2011; Horard *et al*. 2018). Thus, the prevailing view for *de novo* deposition is that ASF1 provides new H3.3 to the HIRA complex, which deposits this variant through the UBN1 subunit. Finally, the HIRA complex is enriched at transcriptionally active regions and can interact with both RNAPII and naked DNA, providing means for H3.3 deposition coupled to transcription (Ray-Gallet *et al*. 2011; Schneiderman *et al*. 2012; Pchelintsev *et al*. 2013). While *de novo* histone deposition could restore nucleosome density, it may not fully restore chromatin profiles and reproduce information carried by old histones. Indeed, newly synthesized histones prior to deposition display a particular set of PMTs distinct from those found into chromatin (Loyola *et al*. 2006). Thus, a key issue is whether and how old H3 histone variants are recycled to provide a template to maintain the pre-existing epigenomic landscape. Lessons from yeast have underlined the importance of histone chaperones to maintain old histones and associated modifications (Nourani *et al*. 2006; Thebault *et al*. 2011; Chen *et al*. 2015; Svensson *et al*. 2015; Jeronimo *et al*. 2019). In mammalian cells, where additional histone variants are present, how the overall choreography of variants and histone modifications is orchestrated *in vivo* during transcription-coupled recycling and by which mechanisms remains unknown.

Here, by exploiting the SNAP-tag system (Keppler *et al*. 2003) for specific *in vivo* labelling to visualize new or old histones in human cells, we address whether and how H3 variants are recycled during transcription. First, we found that transcription results in eviction of old histones, with a stronger impact on H3.3 compared to H3.1. Second, we demonstrate that HIRA is key for the recycling of a large fraction of evicted old H3.3 preventing a major loss of this variant. Surprisingly, this H3.3 recycling mechanism involving the HIRA complex operates through a pathway that differs from the one involved in the deposition of new H3.3, as attested by the fact that neither HIRA trimerization nor UBN1 proved necessary. However, this distinct recycling pathway strictly requires the interaction of HIRA with ASF1 supporting a new role for ASF1 in old H3.3 recycling during transcription. Together, our results reveal that histones are actively recycled during transcription in mammals and identify the mechanism underlying this recycling. Finally, given the central role of the HIRA subunit in both *de novo* deposition and recycling, we discuss how the combination of both can operate through the HIRA trimerization properties to provide means to fine-tune the balance between new and old histones.

## RESULTS

### Transcription causes a global loss of H3.3 in a short time scale

To explore old H3.1 and H3.3 histone variant dynamics, we exploited two HeLa cell lines previously characterized in our laboratory that stably express SNAP-tagged H3.3 or H3.1 (Ray-Gallet *et al*. 2011; Clément *et al*. 2018). Under appropriate experimental conditions, SNAP-tag labelling enables *in vivo* monitoring of total, old or new histones. With this method, we labelled old H3.3- or H3.1-SNAP using a SNAP-compatible tetramethyl-rhodamine fluorophore (TMR) followed by a chase period of 0h, 1h, 2h, 6h, 12h, 24h or 48h (Pulse-Chase) (Figure 1a). We next recorded the TMR signal retained in the nuclei of a population of cells at the different times and used this measure as a proxy to assess histone loss (for details of the methods see Torné *et al*. 2018). In agreement with previous observations (Clément *et al*. 2018), the signal loss for both H3.1- and H3.3-SNAP could not be explained simply by the two-fold dilution expected from cell divisions, occurring every 24 hours in these cells (Figure 1a). Furthermore, signal intensity showed a rapid decrease of 17% for H3.1-SNAP and 36% for H3.3-SNAP in the first two hours, with kinetics that cannot be explained by a single turnover rate (Figure 1a). Considering that the total levels of histones remained stable in our cells, this revealed a rapid exchange for a fraction of H3.1- and H3.3-SNAP histones can be captured within 2h.

**Figure 1.**
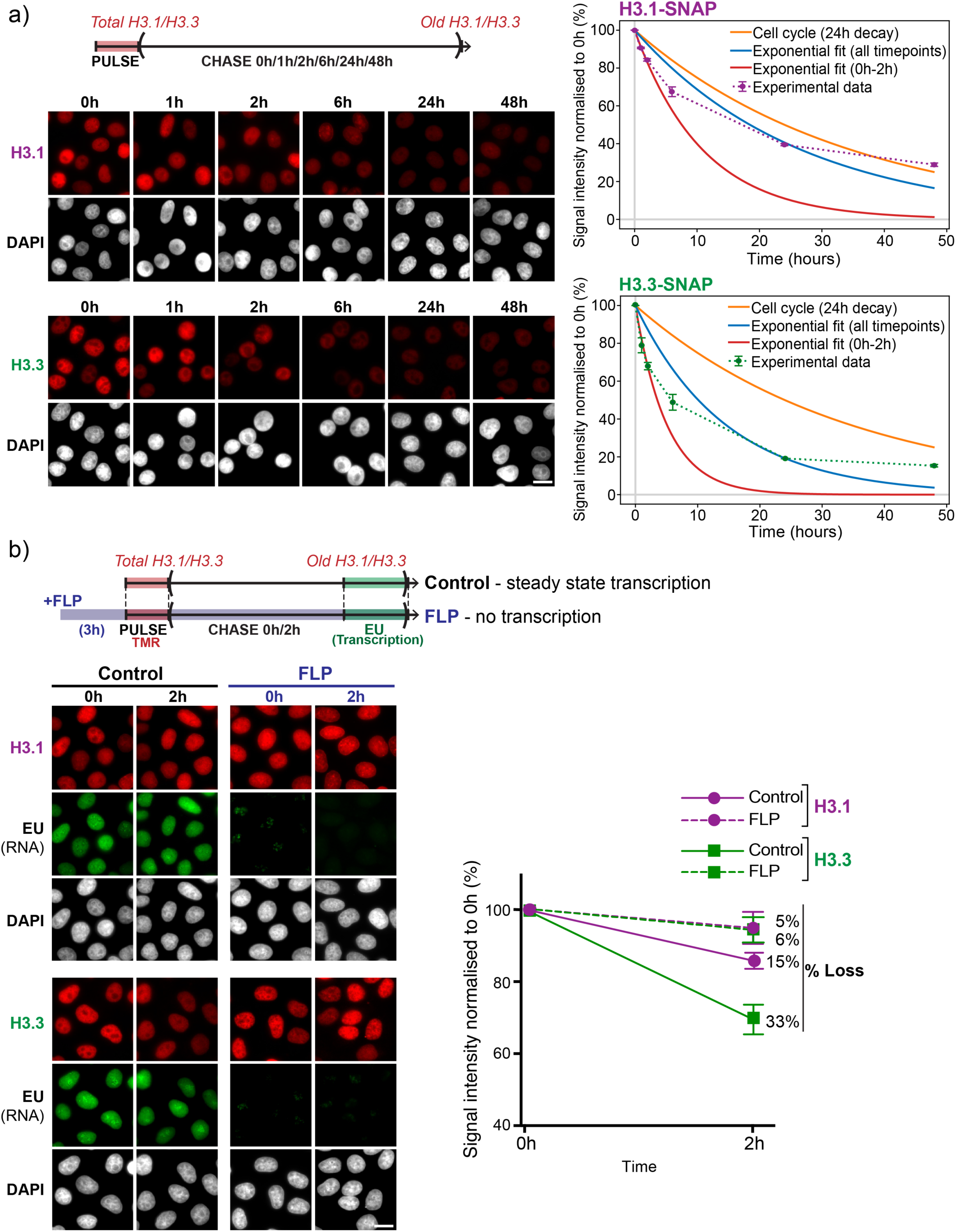
Loss of old H3.3 is dependent on transcriptional activity. Short term old H3.3 loss is dependent on transcriptional activity. **a**) Top: Experimental set-up to track total and old H3.1/H3.3-SNAP (TMR, red) following different chase times (0h to 48h). Bottom-left: representative wide field epifluorescence images of H3.1- and H3.3-SNAP (red), and DAPI (grey). Right: quantification of average TMR signal for H3.1- and H3.3-SNAP, together with its best exponential fit (blue), the exponential fit over 0 to 2h (red) and the expected exponential decay for cell cycle-dependent histone dilution (yellow). **b)** Top: Experimental set-up to track total and old H3.1/H3.3-SNAP (TMR, red) in the presence or absence of the transcription inhibitor Flavopiridol (FLP) for 3 hours prior to TMR pulse and during the chase time. EU labelling (green) marks nascent RNA and is used to confirm the absence of transcription in Flavopiridol-treated cells at every chase time point. Bottom-left: representative wide field epifluorescence images of control and FLP-treated H3.1- or H3.3-SNAP cells after 0h and 2h of TMR chase time. Bottom-right: quantification of average TMR signal for H3.3-(green) and H3.1-SNAP (purple) of untreated (full lines) or FLP-treated (dashed lines) cells, expressed as a percentage of the average value at chase time 0h. For all samples, n>200 nuclei per replicate were analysed. Numbers shown are the average and standard error for independent biological triplicates. Scale bars represent 10 μm.

To further understand the causes for such rapid short-term histone loss, we also labelled newly-synthesized DNA using Ethynil-deoxyuracyl (EdU) at the end of the 0h, 1h or 2h chase times, to distinguish cells undergoing DNA replication. We used EdU signal to identify cells in S-phase (EdU-positive) and outside of S-phase (EdU-negative) (Supplementary Figure 1a) and quantified the TMR signal in these groups of cells to assess histone loss over time. Cells in or outside of S-phase, showed similar H3.1- and H3.3-SNAP loss, indicating that this short-term loss is independent of S-phase progression and showed a more pronounced effect on H3.3 when compared to H3.1.

Given that in mouse ES cells H3.3 is progressively lost at active but not at inactive genes (Deaton *et al*. 2016), we tested whether transcriptional activity could cause H3.3 eviction from chromatin. To this end, we used Flavopiridol – a kinase inhibitor that blocks phosphorylation of NELF, impeding RNAPII release from promoter pausing – to inhibit transcription. We used 5-Ethynyl uridine (EU) to label nascent transcripts in single cells and verified that transcription was reduced to background levels after 3 hours of Flavopiridol treatment (Figure 1b). Under these conditions, we next measured old H3.1- and H3.3-SNAP decay in transcribing versus non-transcribing cells. Without releasing Flavopiridol treatment, we performed a SNAP-tag experiment to label old H3.1- and H3.3-SNAP, and monitored their levels after 0h or 2h of chase time. In non-transcribing cells, H3.1-SNAP signals decayed slowly (5% loss), at slightly lower rates compared to control transcribing cells (15% loss), indicating that transcription arrest modestly alleviates H3.1-SNAP loss (Figure 1b). However, we observed a more dramatic effect for H3.3-SNAP loss, where the loss was reduced to only 6% after 2h of chase, a rate comparable to H3.1-SNAP loss, compared to 33% H3.3-SNAP loss in control transcribing cells. Importantly, blocking transcription with Triptolide – which blocks transcription through inhibition of helicases required for formation of the transcription pre-initiation complex– we observed the same trends (Supplementary Figure 1b). We concluded that within a range of 2h, transcription is the major cause of the loss of old H3.1 and H3.3 variants with a more pronounced effect on H3.3.

### Local dynamics of H3.3 at transcriptionally active domains

Genome-wide analyses previously showed a specific enrichment of H3.3 at transcriptionally active chromatin domains and relative depletion of H3.1 (Goldberg *et al*. 2010; Clément *et al*. 2018). Similarly, at a single cell level for an individual territory in Drosophila polytene chromosomes from salivary glands, the same effect was observed (Schwartz and Ahmad 2005; Schneiderman *et al*. 2012). Yet, the dynamics of exchange of new and old histones has not been directly analysed in human cells. We thus sought to evaluate the spatial relationship between total, new and old H3.3/H3.1 compared to transcriptionally active subnuclear domains in individual human cells exploiting our imaging approach. We thus fluorescently stained our HeLa H3.3/H3.1-SNAP cells: we labelled histone variants with the SNAP-tag method and transcriptionally active forms of RNAPII by immunostaining. Respectively, phosphorylation of Serine 2, 5 and 7 (S2ph, S5ph and S7ph) on the C-terminal tail of RNAPII have been associated with different phases during RNAPII activity, namely promoter pausing (S5ph and S7ph), early elongation (S5ph and S7ph) and late elongation (S2ph and S7ph) (Mayer *et al*. 2010).

As previously described, the active forms of RNAPII appear as discrete foci in the nucleus (Ghamari *et al*. 2013; Cho *et al*. 2018). We focused our analysis on these foci (hereby referred to as transcriptionally active domains). In contrast, as expected from their widespread distribution in the genome, H3.1- and H3.3-SNAP, showed a rather homogeneous distribution in the nucleus (Figure 2). Considering this homogenous distribution the use of common signal colocalization analysis methods to detect changes between different biological conditions was not adapted. To overcome this difficulty, we designed an image analysis method to evaluate the spatial relationship between these two kinds of signals. In this approach, we first segment a primary signal (RNAPII) to identify a set of discrete foci. Next, the secondary signal (histones) is measured within these foci as well as at increasing distances from these foci. The cumulated secondary signal is normalized to total nuclear signal and plotted as relative signal at each distance point from the primary foci. This analysis yields characteristic curves reflecting the spatial enrichment or depletion of the secondary signal compared to the primary signal (Figure 2a and Supplementary Figure 2). We first asked whether H3.3 and H3.1 were enriched at transcriptionally active domains. Our analysis revealed a sharp enrichment of H3.3-SNAP at RNAPIIS7ph foci, corresponding to transcriptionally active domains in single human cells (Figure 2b). In contrast, H3.1-SNAP was depleted from RNAPIIS7ph foci (Figure 2b). The same analysis carried out for RNAPIIS2ph (elongating RNAPII) and RNAPIIS5ph (initiating RNAPII), showed again a characteristic H3.3 enrichment and H3.1 depletion. Thus, this profile applies broadly to transcriptionally active domains throughout promoter pausing, early elongation and late elongation (Supplementary Figure 3a).

**Figure 2.**
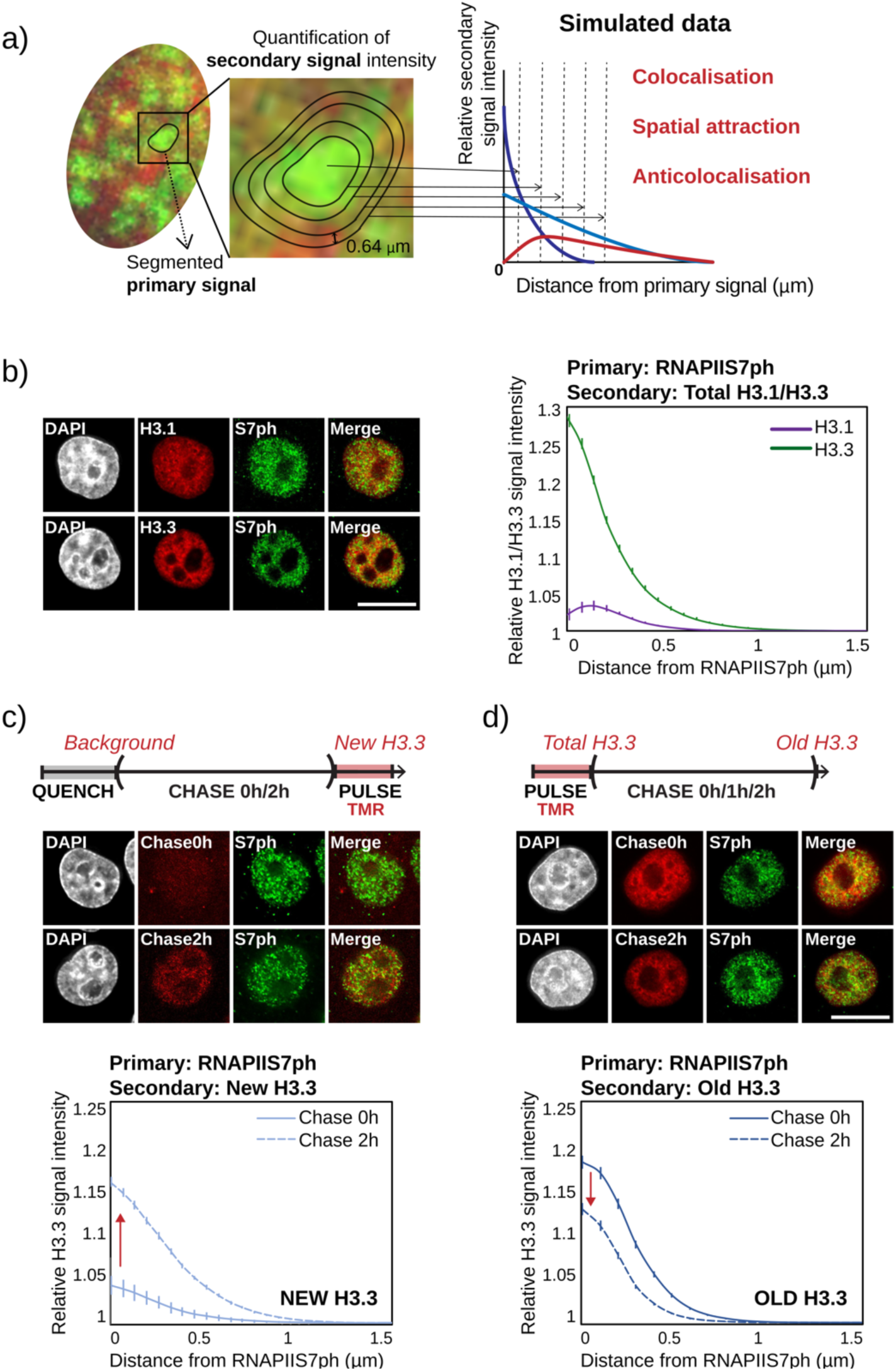
H3.3 is enriched and dynamically exchanged at transcriptionally active domains. **a)** Schematic description of our spatial relationship analysis method. Left: A primary signal (green) is segmented into discrete foci and the intensity of a secondary signal (red) is quantified within and at fixed increasing distances from these foci. Right: results are reported as the background-subtracted average cumulated signal at each distance from primary foci, normalized to total nuclear signal. Characteristic colocalisation, spatial attraction and anticolocalisation curves were obtained with simulated data (see Sup. Fig 2). **b)** Left: representative deconvolved epifluorescence images of cells stained for total H3.1- or H3.3-SNAP (TMR, red), RNAPIIS7ph (green) and DNA (DAPI, grey). Right: spatial relationship analysis showing an enrichment of H3.3 (green) and depletion of H3.1 (purple) at RNAPIIS7ph foci. **c)** Top: Experimental set-up to track new H3.3-SNAP histones synthesized following a BTP quench after 0h (Background) and 2h of chase time (New H3.3). Middle: representative deconvolved epifluorescence images of cells stained for RNAPIIS7ph (green) and new H3.3-SNAP (TMR, red) at 0h and 2h chase time. Bottom: spatial relationship analysis shows enrichment of new H3.3 relative to RNAPIIS7ph at 2h (dashed line) chase time compared to 0h background control, indicating that new H3.3 is preferentially accumulated at these foci. **d)** as in c), except Old H3.3 was tracked at 0h (Total H3.3) and 2h (Old H3.3). Spatial relationship analysis shows depletion of old H3.3 relative to RNAPIIS7ph at 2h (dashed line) chase time compared to 0h total H3.3-SNAP control, indicating that old H3.3 is preferentially lost at these foci. All plots show average and standard error of n>40 cells. Scale bars represent 10 μm.

Further exploiting this approach, we next examined the dynamic exchange of H3.3-SNAP protein localized at these domains. This time, we used the SNAP-tag methodology to label either newly synthesized or old H3.3. We followed old H3.3-SNAP using our Pulse-Chase strategy, as above, and new H3.3-SNAP using the previously described Quench-Chase-Pulse strategy (Ray-Gallet *et al*. 2011). In this labelling scheme to detect newly synthesized histones, old H3.3-SNAP is covalently bound to a fluorescently inert compound, bromothenylpteridine, which prevents TMR binding (Quench). In this scheme, only proteins synthesized after the quench, during the chase time, can bind fluorescent TMR. We measured the spatial distribution of new or old H3.3-SNAP relative to RNAPIIS7ph foci. First, we observed that new H3.3-SNAP became enriched at these transcriptionally active domains within 2h (∼200% gain, Figure 2c). Conversely, old H3.3-SNAP becomes depleted at these domains relative to total H3.3-SNAP within the same chase time of 2h (∼25% loss, Figure 2d). Because our analysis measures enrichment levels relative to total nuclear signal, these results indicate that new and old H3.3 are respectively gained and lost preferentially at transcriptionally active domains, compared to the rest of the nucleus. We observed the same profiles when using RNAPIIS2ph and RNAPIIS5ph as primary signals, further validating our observations (Supplementary Figure 3b and 3c). These results indicate that H3.3 is enriched at transcriptionally active domains in single cells, where old H3.3 is frequently replaced by new H3.3. Together, our results show a preferential exchange of H3.3 at transcriptionally active domains, in a transcription-dependent manner.

### Retention of old H3.3 at transcriptionally active domains requires the chaperone HIRA

While H3.3-SNAP loss was remarkable (36% in two hours), we wondered if the significant fraction that remained (64%), was being actively retained. The next question was thus whether and how this could be achieved. Using a candidate approach, we focused on the H3.3 chaperone HIRA. We performed efficient knockdown of HIRA in our SNAP-tag cells, as previously described (Ray-Gallet *et al*. 2011) and labelled new or old H3.1- and H3.3-SNAP to track their fate during 0h, 1h or 2h of chase time. First, we confirmed that HIRA knockdown dramatically impacts new H3.3 deposition, without affecting H3.1 (Ray-Gallet *et al*. 2011) (see also Figure 5a and Supplementary Figure 5). Next, when following old histones upon HIRA knockdown, strikingly, we observed that the loss of H3.3-SNAP was more dramatic, reaching an average 62% loss in 2h, compared to 34% loss in control conditions (Figure 3a). Interestingly, old H3.1-SNAP loss was mildly alleviated in knockdown cells (4% loss), compared to 15% loss in control cells, possibly reflecting a compensation of the massive H3.3-SNAP loss. These results demonstrate that HIRA is required not only for deposition of new H3.3, but also for retention of an important fraction of old H3.3.

**Figure 3.**
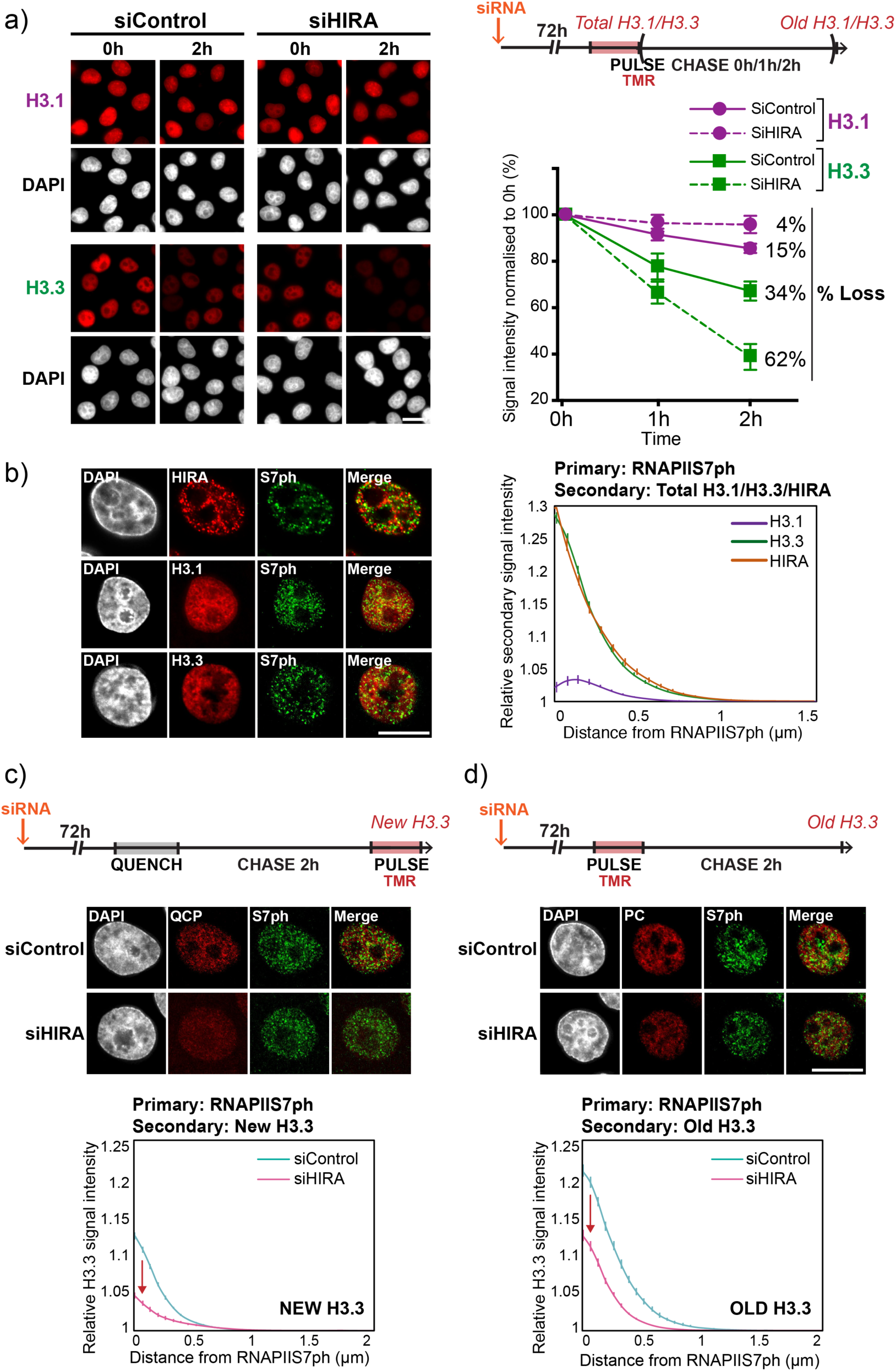
HIRA is required for both deposition of new H3.3 and retention of old H3.3 at transcriptionally active domains. **a)** Top-right: Experimental strategy to track old histones in cells treated with control or HIRA-targeting siRNA 72h prior to SNAP-tag labelling. Left: representative wide field epifluorescence images of cells stained for H3.1- or H3.3-SNAP following control (siControl) or HIRA-targeting (siHIRA) siRNA treatment, and after 0h and 2h of TMR chase time. Bottom-right: quantification of average nuclear TMR signal for H3.1 (purple) and H3.3 (green), in control (siControl, full lines) and HIRA knockdown (siHIRA, dashed lines) cells, expressed as a percentage of the average value at chase time 0h. The average percentage of loss at 2h chase time is indicated for each sample, indicating that H3.3 is more rapidly lost upon HIRA knockdown, while H3.1 loss is alleviated in these conditions. For each sample, n>200 cells were analysed. Plots show the average and standard error of independent biological triplicates. **b)** Left: representative deconvolved epifluorescence images for cells stained for total H3.1- or H3.3-SNAP (TMR, red) or using a HIRA antibody (red), together with RNAPIIS7ph (green) and DNA (DAPI, white). Right: spatial relationship analysis plots showing enrichment of HIRA (orange) and H3.3 (green) but not H3.1 (purple) at RNAPIIS7ph foci. **c)** Top: Experimental strategy to track new H3.3 upon HIRA knockdown. Middle: representative deconvolved epifluorescence images of cells stained for new H3.3 (TMR, red), RNAPIIS7ph (green) and DNA (DAPI, grey). Bottom: spatial relationship analysis showing preferential depletion of new H3.3 at RNAPIIS7ph foci in the absence of HIRA. **d)** As in c), except old H3.3 was tracked, showing that old H3.3 is also preferentially depleted at RNAPIIS7ph foci in the absence of HIRA. For spatial relationship analysis, numbers are averages from n>40 cells. Scale bars represent 10µm.

Following our observation of a dynamic exchange of H3.3 at transcriptionally active domains, we further sought to directly test a role for HIRA in deposition of new H3.3 and retention of old H3.3 locally at these domains. We first performed immunostaining of HIRA together with phosphorylated RNAPII to evaluate their spatial relationship. We found a sharp enrichment of HIRA at RNAPIIS7ph foci (Figure 3b), as well as foci of RNAPIIS2ph and RNAPIIS5ph forms (Supplementary Figure 4a). This chaperone is thus specifically enriched at transcriptionally active domains in single cells, consistent with previous genome-wide data (Pchelintsev *et al*. 2013). To evaluate its role to guide H3.3 dynamics at these sites, we further performed HIRA knockdown in SNAP-tag cells and labelled new or old H3.3-SNAP together with RNAPIIS7ph. Consistent with other reports, we noticed a modest but significant (22%) impact of HIRA depletion on global transcriptional activity, with Polymerases accumulating on chromatin while transcription itself is perturbed (Supplementary Figure 5) (Maze *et al*. 2015; Zhang *et al*. 2017). We further applied our imaging analysis method to evaluate the fate of new and old H3.3-SNAP at transcriptionally active domains. Upon HIRA knockdown, we measured a depletion of both new and old H3.3-SNAP at these domains, compared to control knockdown conditions (±60% loss and ±50% loss respectively). Again, since our analysis measures enrichment levels relative to total nuclear signal, this indicates that depletion of new and old H3.3-SNAP occurs preferentially at transcriptionally active domains (Figure 3c, 3d and Supplementary Figure 4). Together, our results show that HIRA is required for both deposition of new and retention of old H3.3, with an exacerbated effect at transcriptionally active domains.

To further test whether this HIRA requirement is linked to transcription, we inhibited transcription using Flavopiridol (see Figure 1). As above, we observed a higher retention of old H3.3-SNAP on chromatin in the absence of transcription with only 10% loss (Figure 4). In this scenario, HIRA knockdown had no longer an effect on this H3.3-SNAP retention (8% loss). We concluded that in absence of transcriptional activity, HIRA is not required to re-deposit the H3.3 variant. We confirmed these results using an independent method for transcription inhibition using Triptolide (Supplementary Figure 6). Together, our results demonstrate that HIRA is essential to recycle a fraction of old H3.3 evicted by transcriptional activity.

**Figure 4.**
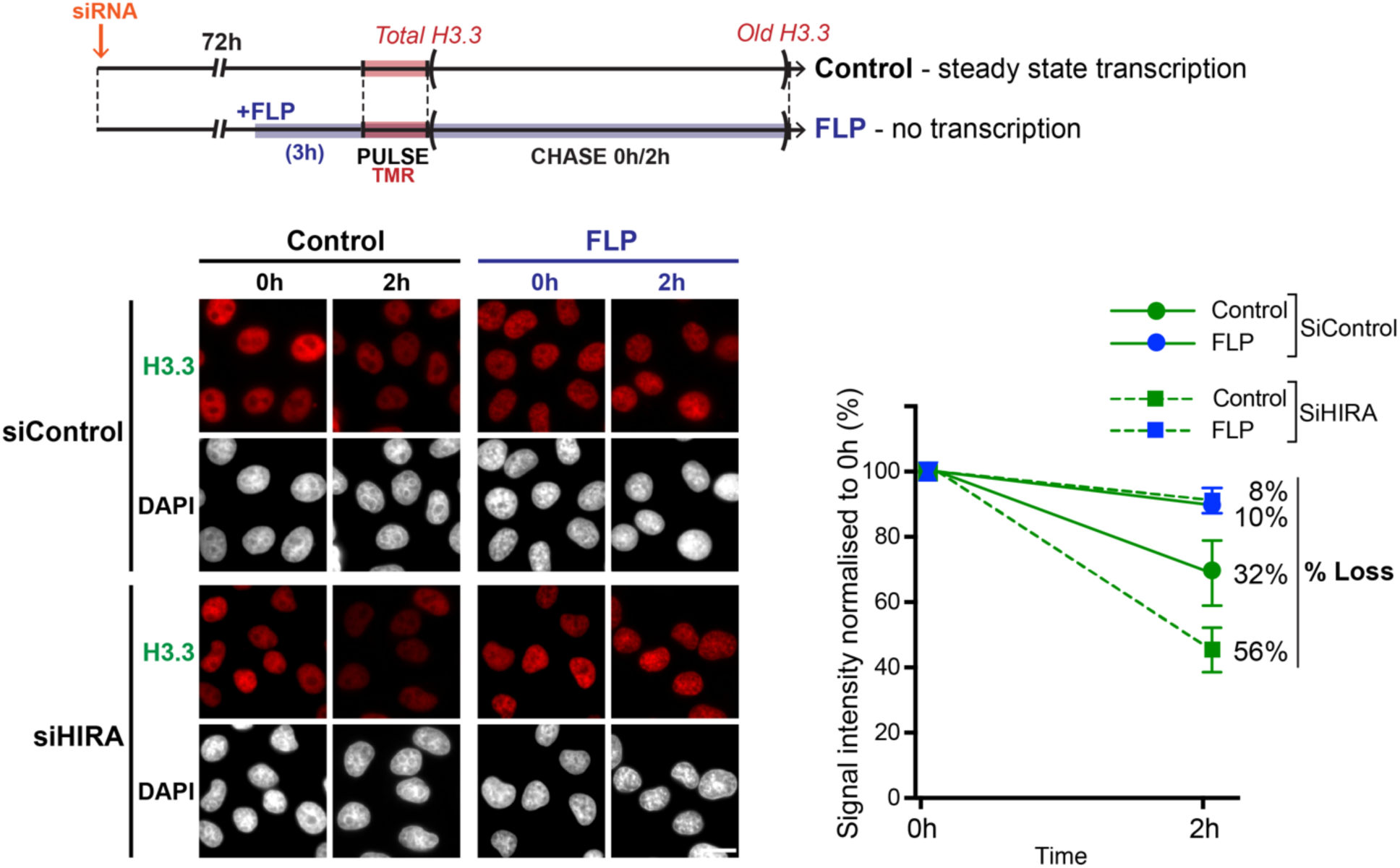
Transcription is required to reveal the HIRA dependent H3.3 recycling. Top: experimental strategy to track old H3.3 during steady state transcription and in cells exposed to FLP until transcription was fully arrested (as in Figure 1b). Bottom-left: representative images of total (0h) or old (2h) H3.3-SNAP (TMR, red), in control conditions or during FLP treatment and 72h following knockdown using Control or HIRA-targeting siRNA. Bottom-right: quantification of TMR signal for control (full lines) and HIRA knockdown (dashed lines), untreated (green) and FLP-treated (blue) cells indicate that, in transcriptionally-arrested cells, absence of HIRA has no effect on old H3.3 dynamics. For all samples, n>200 cells. Plots show averages and errors from independent biological triplicates. Scale bars represent 10µm.

### *De novo* deposition and recycling of H3.3 by HIRA occurs through distinct pathways

To define how HIRA specifically handles new and old H3.3, we next investigated the roles of known HIRA chaperone partners: UBN1, CABIN1 and ASF1. As described above, UBN1 is the subunit of the HIRA complex recently described as a key interacting partner for new H3.3 deposition (Ricketts *et al*. 2015; 2019), while ASF1 is rather an upstream chaperone that supplies histones to the HIRA complex. We performed individual knockdowns of these factors and used the SNAP-tag strategy to track new and old H3.3 dynamics. We confirmed that knockdown of HIRA and UBN1 impacted new H3.3-SNAP deposition, while CABIN1 knockdown had no effect (Figure 5a), as described (Ray-Gallet *et al*. 2011). Simultaneous knockdown of both ASF1a and ASF1b isoforms moderately impaired H3.3-SNAP deposition, to a lesser extent than HIRA or UBN1 knockdown. Interestingly, double knockdown of ASF1a/b also led to increased old H3.3-SNAP loss compared to control cells, with 51% loss over 2h (Figure 5b). This effect is milder than that of HIRA knockdown (62%), possibly because transcriptional activity itself is reduced (41%), as assessed with EU labelling in ASF1a/b double knockdown cells (Supplementary Figure 5b). Surprisingly, knockdown of UBN1 had no effect on old H3.3-SNAP retention (Figure 5b) or nascent transcription (Supplementary Figure 5b). This indicated that HIRA is capable of recycling H3.3 independently of its partner UBN1. Thus, HIRA operates through two distinct pathways for *de novo* deposition and recycling of H3.3.

**Figure 5.**
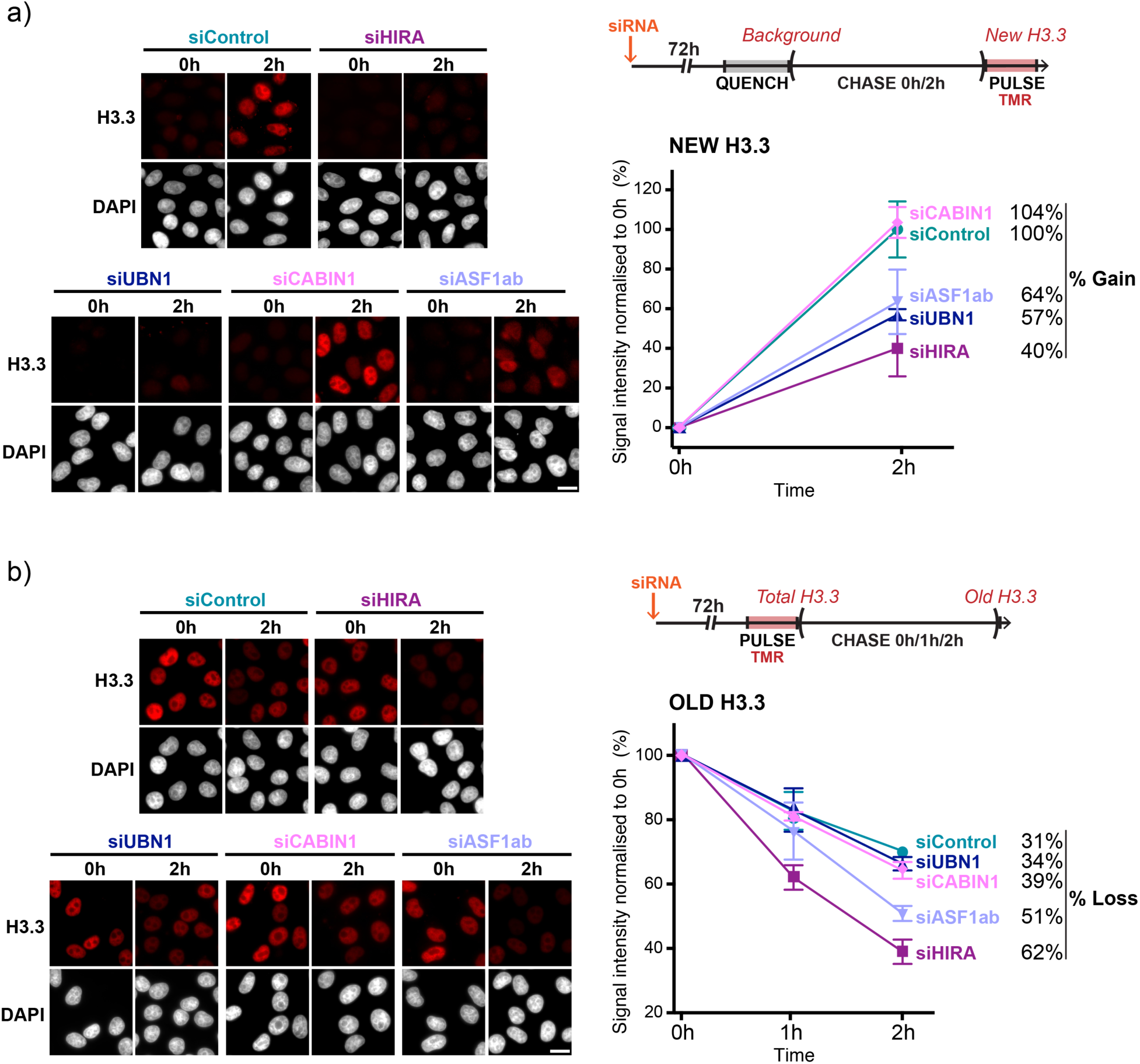
Different partnerships for HIRA in new H3.3 deposition versus recycling. **a)** Top-right: experimental scheme to track new H3.3-SNAP in cells treated with siRNA for 72h. Left: representative wide field epifluorescence images of cells stained for new H3.3-SNAP after 0h (Background) or 2h of chase time (New H3.3) (TMR, red), and DNA (DAPI, grey), 72h after knockdown of HIRA, UBN1, CABIN1 or ASF1 isoforms a+b or using a control siRNA. Bottom-right: quantification of total nuclear TMR signal for each knockdown condition expressed as a percentage of H3.3 gain in 2h relative to the control condition. The results indicate that new H3.3 deposition requires HIRA, UBN1 and ASF1 but not CABIN1. **b)** as in a), except old H3.3-SNAP was tracked after 0h (Total H3.3), 1h or 2h (Old H3.3) of chase time, indicating that old H3.3 recycling requires HIRA and ASF1 but not UBN1 or CABIN1. For all samples, n>200 cells were imaged per replicate. All plots show averages and standard errors for two (panel a) or three (panel b) biological replicates.

To further explore the underlying mechanism, we used several HIRA-YFP constructs in which single amino-acid substitutions have been introduced in order to disrupt the interaction with particular partners (Figure 6a). The R227K mutant in the WD40 domain of HIRA disrupts its interaction with UBN1 (Loppin *et al*. 2005; Banumathy *et al*. 2009). The I461D mutant in the conserved B-domain of HIRA prevents its interaction with ASF1 (Tang *et al*. 2006). We also recently described the W799A-D800A mutant, containing a double amino acid substitution, which prevents both trimerization of the HIRA protein and its interaction with CABIN1 (Ray-Gallet *et al*. 2018). With each of these HIRA mutants, we could thus test their capacity to rescue the H3.3 *de novo* deposition and recycling defects caused upon HIRA knockdown. To track and selectively analyse transfected cells, all HIRA proteins were tagged with YFP. We verified that all transgenic proteins in the transfected cells showed a comparable expression level (Supplementary Figure 7a-b). Wild-type HIRA could readily rescue new deposition of H3.3-SNAP (Figure 6b-c). In contrast, the HIRA mutants defective for UBN1 interaction (HIRA-R227K-YFP) or HIRA trimerization (HIRA-W799A-D800A-YFP) did not alleviate the defect in new H3.3-SNAP deposition. This observation confirmed that HIRA requires both interaction with UBN1, as well as the ability to trimerize in order to efficiently deposit new H3.3 (Ray-Gallet *et al*. 2018). The HIRA mutant that impairs ASF1 interaction (HIRA-I46D-YFP), while less efficient than wild type, could partially rescue the H3.3-SNAP deposition. This observation suggests a direct transfer of new H3.3 to UBN1 by exploiting a possible bypass of the HIRA-ASF1 interaction. We next focused on the capacity of these HIRA mutants to rescue old H3.3-SNAP loss (Figure 6d-e). In contrast to wild-type HIRA, this same mutant (HIRA-I46D-YFP) unable to interact with ASF1 could not at all rescue the loss of old H3.3-SNAP. Together, they support an absolute requirement for HIRA to interact with ASF1 to ensure the recycling of old H3.3. Strikingly, targeting UBN1, the HIRA-R227K-YFP mutant (UBN1 interaction defective) fully rescued the old H3.3-SNAP loss associated with HIRA knockdown. This result confirmed that HIRA interaction with UBN1 is dispensable for H3.3 recycling. Finally, the HIRA-W799A-D800A-YFP mutant (impaired for trimerization) also rescued old H3.3-SNAP loss, showing that HIRA does not need to oligomerize to recycle H3.3. This latter observation also confirmed that CABIN1 was not required either in this setting. Together, these results (summarized in Figure 6f) are consistent with our knockdown experiments and bring into light the existence of two distinct pathways for HIRA handling respectively new and old H3.3. New H3.3 deposition requires both HIRA trimerization as well as its interaction with UBN1 and to a lesser extent with ASF1. In contrast, old H3.3 is handled only by ASF1 and HIRA, and does not require either UBN1 or HIRA trimerization.

**Figure 6.**
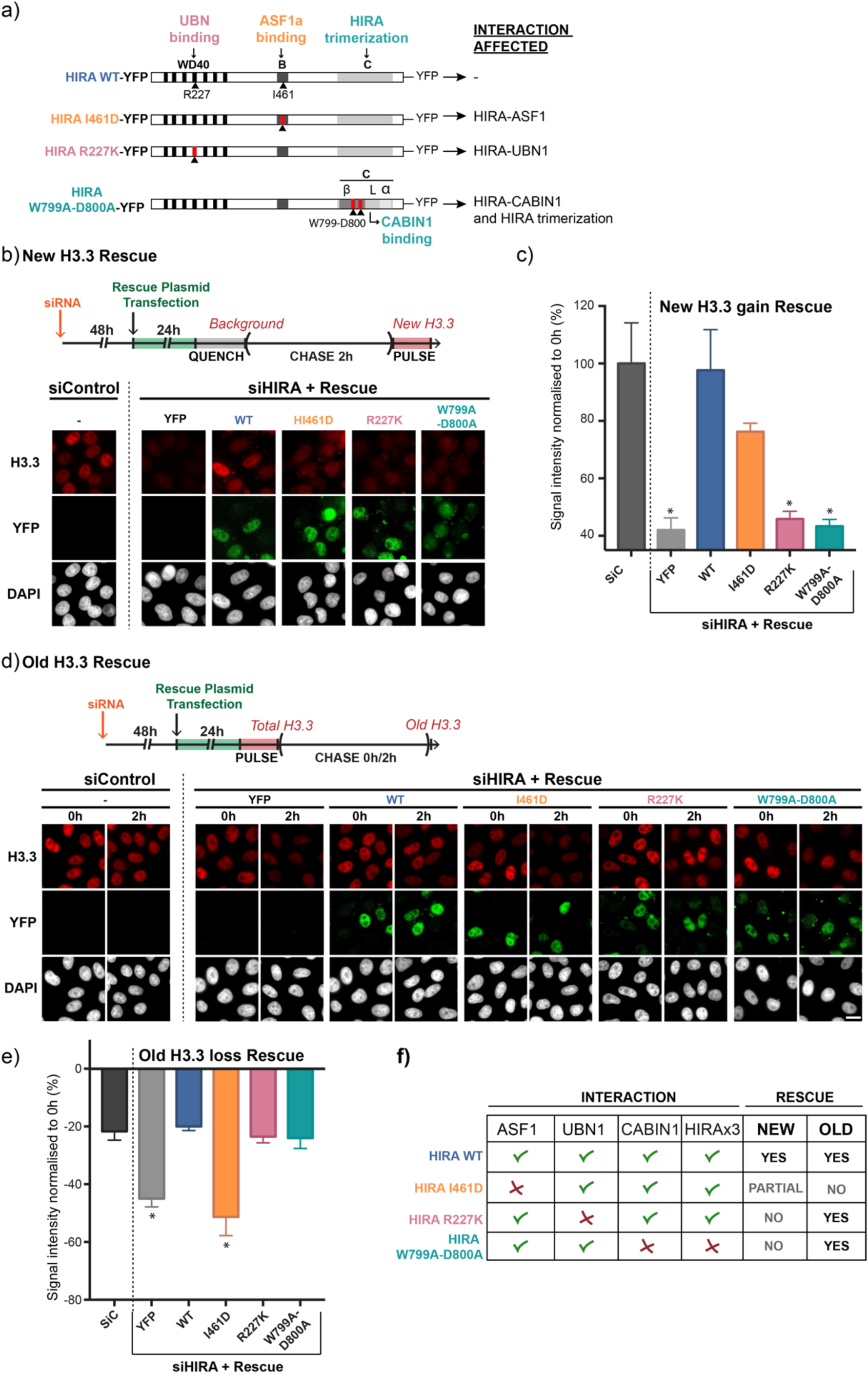
The interaction domain between HIRA and ASF1 is required for old H3.3 recycling. **a)** Scheme depicting mutated HIRA-YFP transgenic proteins used for rescue experiments. Functional domains required for UBN1 interaction (WD40), ASF1 interaction (B-domain) and trimerization (C domain) are shown. Red indicates substituted amino acids. **b)** Top: Experimental strategy to rescue the effect of HIRA knockdown on new H3.3-deposition using HIRA-YFP transgenes. Bottom: Representative images for rescue experiment of impaired new H3.3 deposition with the different transgenic proteins. Cells were imaged for new H3.3-SNAP (TMR, red), as well as YFP (green), and DNA (DAPI, grey). YFP protein alone is undetectable in triton-extracted cells, but readily visible in all HIRA-YFP constructs. **c)** Quantification of total nuclear TMR signal from all conditions expressed as a percentage of new H3.3 gain relative to the siControl sample, indicating that HIRA trimerization and its interaction with UBN1 are required for new H3.3 deposition while ASF1 interaction can be partially bypassed. **d)** as in b), except the effect on old H3.3-SNAP loss is visualized. **e)** Quantification of total nuclear TMR signal from all conditions expressed as a percentage of total H3.3 in siControl sample, indicating that HIRA interaction with ASF1 is essential to recycle old H3.3, while its trimerization or interaction with UBN1 are dispensable. **f)** Summary table for all new and old H3.3 recue experiments results. For all samples, n>200 cells were analysed. Plots show averages and standard errors for two (panel c) or three (panel e) biological replicates (* indicates p-value < 0.05 on a standard t-test). Scale bars represent 10µm.

### HIRA serves as a molecular hub for transcription-coupled deposition of new and old histones

To gain insights into how old histones at transcriptionally active domains are handled, we decided to follow a histone mark associated with active transcription onto chromatin. H3K36me3 is imposed by the methyltransferase Setd2, which travels with RNAPII during transcription (Yoh, Lucas, and Jones 2008; Edmunds, Mahadevan, and mahadevan 2008). Marking transcriptionally active gene bodies (Bannister *et al*. 2005;) and prominently detected in chromatin (Loyola *et al*. 2006), this mark thus represents a proxy for old, nucleosomal histones. Indeed, H3K36me3 is observed by Western blot mainly in chromatin fraction and faintly detected in nuclear extract (Figure 7a). We thus performed an immunoprecipitation with antibodies against H3K36me3 to identify its binding partners using nuclear extracts in which the HIRA complex is enriched. RNAPII co-immunoprecipitated with H3K36me3, as well as the chaperones HIRA and ASF1. In contrast neither the H3.3 chaperone DAXX or the p60 subunit of the replicative H3.1 chaperone CAF-1 were retrieved (Figure 7a). This result further indicates that both HIRA and ASF1 are involved in handling old histones after their transcription-coupled eviction from chromatin.

**Figure 7:**
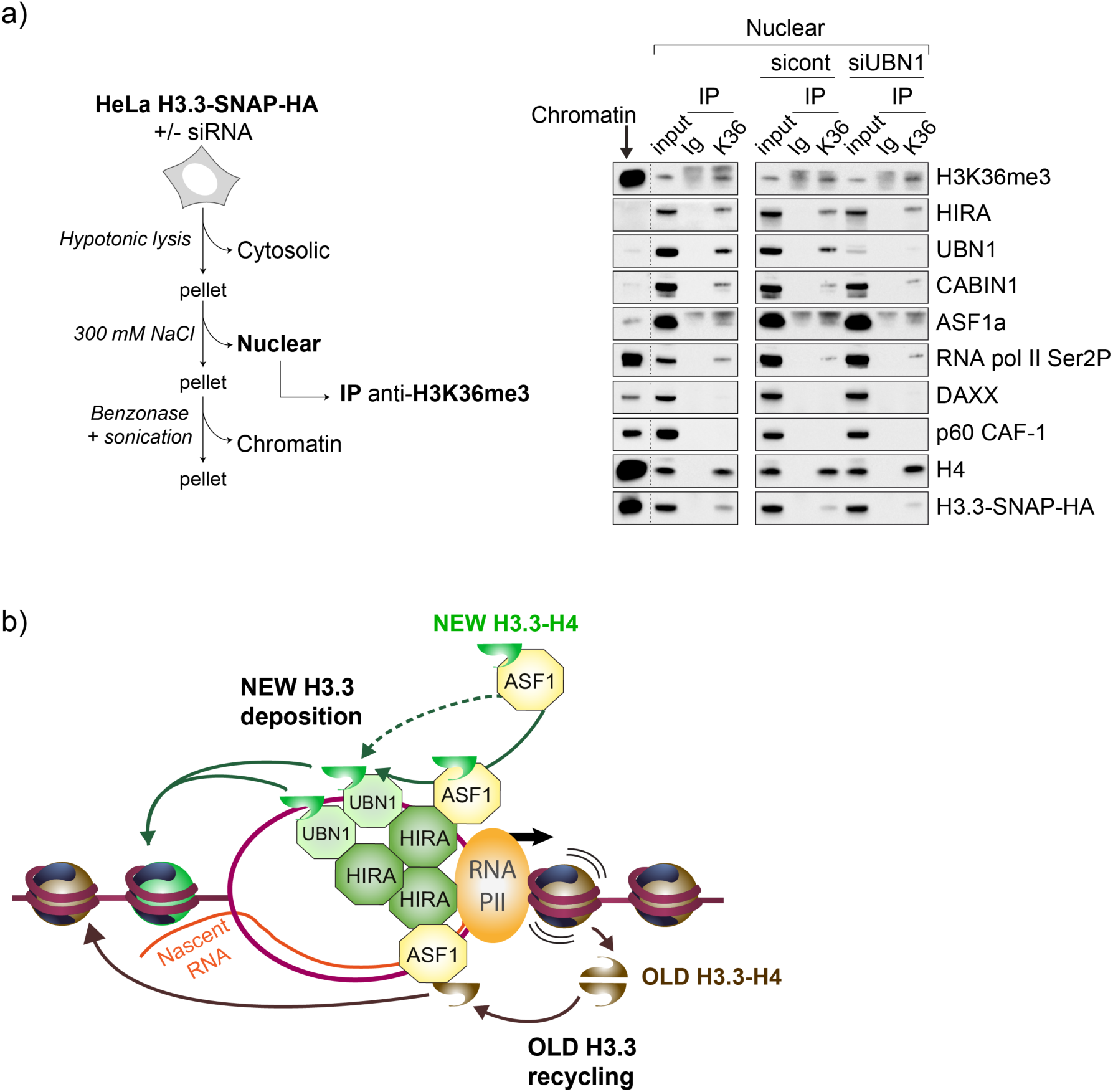
The HIRA complex coordinates deposition of new H3.3 via UBN1 and recycling of old H3.3 via ASF1. **a)** Left: Scheme showing the steps for the preparation of nuclear and chromatin fractions from HeLa cells. Right: western blot analysis of whole chromatin fraction extracts and anti-H3K36me3 (K36) or control rabbit IgG (Ig) immunoprecipitates from nuclear fractions prepared from HeLa H3.3-SNAP-HA cells untreated or treated with control or UBN1-targeting siRNA. Input corresponds to 6% of nuclear extract (18 µg) used for each immunoprecipitation. **b)** Proposed model: A HIRA trimer serves as a platform to coordinate deposition of new and recycling of old H3.3. New deposition relies on HIRA in partnership with UBN1. ASF1 interacting with HIRA hands new H3.3 to UBN1 (represented here as a dimer, as proposed in Ricketts *et al*., 2019 (Ricketts *et al*. 2019); note however that this stoichiometry was not addressed in our study). However HIRA-ASF1 interaction can be bypassed and new H3.3 directly transferred from ASF1 to UBN1 (dashed arrow). In contrast, old H3.3 recycling relies on HIRA independently of UBN1. In this case the interaction of HIRA with ASF1 is absolutely necessary for old H3.3 recycling. For simplicity, the subunit CABIN1 is not represented.

Importantly, UBN1 also co-immunoprecipitated with the H3K36me3 mark. Since this subunit from the HIRA complex is not strictly required for old histone recycling, but is necessary for new deposition, it was critical to verify if depletion of UBN1 would affect H3K36me3 interaction with HIRA and ASF1. We thus performed knockdown for UBN1, and again carried out an immunoprecipitation of H3K36me3 with its partners. We found that UBN1 knockdown did not affect H3K36me3 interaction with HIRA or ASF1 (Figure 7a). These results indicated that HIRA and ASF1 interact with old histones independently of UBN1.

## DISCUSSION

Our work provides a new view on the fate of H3 histone variants-old and new-during transcription. We first establish that transcription leads to a major loss of both old H3.1 and H3.3 variants, with a prominent effect on H3.3. Second, we identify a key mechanism to ensure a significant level of recycling of old H3.3, operating along with new deposition. While both recycling and new deposition pathways exploit the histone chaperone HIRA, we find that the choice between them relies on HIRA partners. We discuss how the histone chaperone HIRA can coordinate H3.3 recycling with new deposition during transcription through interaction with multiple partners and thereby play a pivotal role for the maintenance of chromatin integrity in transcribed regions.

### A dynamic exchange between old and new H3.3 at transcriptionally active domains

In a previous study, we monitored the loss of H3.1- and H3.3-SNAP over the time scale of several cell divisions and found that their kinetics of decay did not fit with a simple exponential curve as expected for a model in which a two-fold dilution occurs at each cell division (Clément *et al*. 2018). A general trend was a faster loss. Here, we monitored the loss of H3.1- and H3.3-SNAP during a shorter time course and our data further underline a short-term loss concerning a fraction of H3.3 and H3.1 that cannot be simply explained by dilution due to cell division. Our results show a replication-independent widespread loss of old H3.3 (36% of SNAP signal decrease over 2 hours, consistent with previous reports (Deaton *et al*. 2016)) and to a lesser extent old H3.1 (17% in 2 hours). Importantly, signals corresponding to both H3.3 and H3.1 total levels are stable in our conditions, indicating that new histone deposition could entirely compensate these losses.

To explore these dynamics of old and new variants within the nuclear space and in relation with transcription, we first examined the relative subnuclear localization of active RNAPII, HIRA and H3.3. Our results show a spatial proximity between them at the scale of nuclear sub-compartments of ∼300nm in diameter, relatively large domains likely to host multiple genes and intergenic DNA. Consistently, we previously described by ChIP-seq a genome-wide profile for H3.3 where this variant covers megabase-scale domains, coinciding with early-replicating gene-rich genomic regions (Clément *et al*. 2018). Furthermore, using super-resolution microscopy, we previously showed how H3.3 could form small clusters, of ∼100nm in diameter (Clément *et al*. 2018), consistent with other reports (Ricci *et al*. 2015; Nozaki *et al*. 2017; Otterstrom *et al*. 2019) and indicating that larger transcriptionally active sub-compartments likely cover an ensemble of smaller histone clusters. Multiple recent studies have proposed that transcriptional domains may feature phase separation properties with intrinsic dynamics tightly linked to transcriptional activity (Cho *et al*. 2018; Sabari *et al*. 2018; Lu *et al*. 2018; Boehning *et al*. 2018; Shaban *et al*. 2018; Guo *et al*. 2019; Nagashima *et al*. 2019). Because of these properties, it is possible that evicted histones could become re-deposited on chromatin close to their original position, or rather be redistributed elsewhere yet contained within these nuclear sub-compartments. These precise dynamics as well as the role of chaperones in their control need to be further elucidated to fully characterize the precise preservation of epigenetic information at different scales.

### A critical role for HIRA in handling both new and old H3.3 during transcription

Our results further show that, in the absence of HIRA, old H3.3 loss is dramatically increased (Figure 3a), indicating that a significant fraction of evicted H3.3 can be recycled by a mechanism involving this chaperone. In addition, when transcription is arrested, the absence of HIRA does not have any impact in H3.3 loss (Figure 4). Thus, when histones within chromatin are no longer challenged by transcription, the requirement for HIRA is essentially abrogated. Consistently, by microscopy analysis accessing these dynamics at a single cell level, we could show that in the absence of HIRA, both new and old H3.3 are specifically lost at transcriptionally active domains (Figure 3c-d). In these conditions, H3.3 homeostasis is thus no longer ensured and this variant is progressively lost from chromatin. Our results thus demonstrate that H3.3 is recycled during transcription and uncover a novel role for the chaperone HIRA in this process, adding to its already known role in *de novo* deposition. While our data place HIRA as an important actor in this recycling, we do not exclude that it could operate in combination with other nucleosome retention mechanisms yet unexplored in mammals. Indeed, in yeast, although there is no distinct H3 variants, histone chaperones including FACT, Spt2 and Spt6 have been reported to have an analogous role during transcription (Nourani *et al*. 2006; Thebault *et al*. 2011; Chen *et al*. 2015; Svensson *et al*. 2015; Jeronimo *et al*. Robert 2019). In addition, there is the intriguing possibility that RNAPII could bypass nucleosomes by orchestrating their 3’ to 5’ transfer, as highlighted by recent crystallography studies (Farnung *et al*. 2018; Vos *et al*. 2018; Kujirai *et al*. 2018). Future work will be needed to examine whether and how these mechanisms could operate *in vivo* in mammalian cells and how they could act in combination with HIRA and the distinct H3 variants. Most importantly, our findings explain how, despite the disruptive nature of the transcriptional process, a histone homeostasis mechanism orchestrated by HIRA, in concert with mechanisms ensuring the spreading of histone marks, could ensure that the epigenomic landscape remains stable at transcribing regions.

### New and old H3.3 are handled by distinct pathways

One could have assumed that a simple HIRA-mediated pathway could indistinctly handle both old and new histones to be deposited. Yet, a surprising finding in our study is that H3.3 recycling involves HIRA interacting with its partner ASF1, but does not require UBN1 nor HIRA homotrimerization (Figures 5-6). In mammals, another Ubinuclein exists, UBN2, that is also able to interact with HIRA (Banumathy *et al*. 2009). However, we discard a potential compensation by UBN2 in the absence of UBN1 as UBN2 cannot interact with the HIRA-R227K mutant (Banumathy *et al*. 2009), which readily rescued old H3.3 loss. Yet, UBN1 and the HIRA homo-trimerization are absolutely necessary for the *de novo* deposition of H3.3. Thus, we can discard a model whereby old H3.3 evicted from chromatin could be treated as new H3.3 after joining the soluble pool of histones for re-deposition. Instead, new H3.3 is guided to chromatin by a dedicated pathway depending on ASF1, UBN1 and a HIRA trimer, while old H3.3 is handed over by ASF1 to HIRA, without requiring HIRA trimerization and UBN1 interaction (Figure 7b). The way ASF1 handles distinctly new and old H3.3-H4 is remarkable. Indeed, for the new H3.3 deposition, the partial rescue by the HIRA-I461D mutant, that cannot interact with ASF1, suggests that ASF1 could transfer H3.3-H4 directly to UBN1 bypassing a need to interact with HIRA (Horard *et al*. 2018). In contrast, this HIRA-I461D mutant fails completely to rescue old H3.3 recycling indicating the absolute requirement of this interaction with ASF1 for handling old H3.3. For *de novo* deposition, given the fact that newly synthesized H3-H4 have been isolated as dimers (Tagami *et al*. 2004), the fact that the UBN1 unit can dimerise offered an attractive means to ensure the formation of a new (H3.3-H4)_2_ tetramer prior/or immediately at the time of incorporation into chromatin (Ricketts *et al*. 2019). One can envisage though that capturing parental histones may require different properties that only ASF1 would have. Notably, there are histone PTMs which are specific for new or old (nucleosomal) histones (Loyola *et al*. 2006). In human cells, in the absence of ASF1, by super resolution microscopy, we could observe a redistribution of histone PTMs, including the old histones PTM H3K36me3, (Clément *et al*. 2018). In this regard, our experiments linking H3K36me3 to the HIRA-ASF1 recycling pathway are particularly enlightening. Future work should explore how, beyond H3K36me3, other PTMs could be preferentially recycled or lost during transcription and reveal the molecular details by which UBN1 and ASF1 could distinguish new and old histones.

The placement and structure of HIRA in transcription shows an interesting parallel to the role of the replisome component Cohesion establishment factor 4/Acidic nucleoplasmic DNA-binding protein-1 (Ctf4/AND-1) (Simon *et al*. 2014; Guan *et al*. 2017). This protein shares a number of structural similarities with HIRA, in particular a similar homo-trimeric structure (Ray-Gallet *et al*. 2018), it associates with the replication fork and is involved in the replication-coupled histone recycling (Gan *et al*. 2018). It is thus tempting to speculate that as Ctf4/AND-1, in replication, HIRA may act as a scaffold to recruit different partners for histone handling during transcription. This view places HIRA as a central player in a mechanism ensuring a balance between the use of new and old histones and also other partners related to DNA/RNA metabolism. In the latter case, a role for HIRA trimerization and its association with UBN1 and ASF1 in the context of transcription restart after DNA damage would deserve to be explored (Adam *et al*. 2013). Furthermore, regulating the balance of new versus old histone deposition may actually prove important when the transcription machinery encounters potential blockade (Gregersen and Svejstrup 2018) to assist dynamics associated with complex chromatin disruption and loss of parental histones.

In conclusion, our study reveals that the histone variant H3.3 is recycled during the process of transcription by HIRA, which acts as a hub to coordinate new H3.3 deposition and old H3.3 re-deposition in collaboration with different partners. These findings highlight the importance of retaining old histones at transcriptionally active regions, a new histone homeostasis pathway to maintain chromatin integrity and pre-existing epigenetic information.

## MATERIALS AND METHODS

### New and Old H3.3- and H3.1-SNAP labelling *in vivo*

We used cell lines stably expressing H3.3-SNAP-3xHA or H3.1-SNAP-3xHA in HeLa cells, previously used and characterized in our lab (Ray-Gallet *et al*. 2011). To track old histones, we followed the Pulse/Chase strategy. We incubated our cells in complete medium containing 2 μM of SNAP-Cell TMR-Star (New England Biolabs) during 20 min to label all pre-existing available SNAP-tag (Pulse). After rinsing twice with phosphate-buffered saline (PBS) we re-incubated the cells in complete medium for 30 min to allow excess SNAP-Cell TMR-Star to diffuse out. We next incubated the cells with complete medium during 0h (i.e Total H3.1- or H3.3-SNAP), 1h or 2h (Chase). To track new histones, we followed the Quench/Chase/Pulse strategy. We incubated cells in complete medium containing 10 μM of SNAP-Cell Block (New England Biolabs) to block all available pre-existing SNAP-tag (Quench), followed by two PBS washes and 30 min of incubation in complete medium to allow the SNAP-Cell Block to diffuse out. We next incubated cells in complete medium for a 0h (i.e background levels), 1h or 2h period (Chase), then performed TMR-Star labelling (Pulse) as described above. If nascent DNA or RNA labelling was required, cells were incubated with 10 μM of EdU or EU respectively during the last 30 min of the experimental pipeline. At least three independent experiments were performed for each condition.

### Extraction and fixation followed by EdU or EU detection

We performed a pre-extraction by incubating cells at room temperature for 5 min in 0.5% Triton in CSK buffer (10 mM PIPES (pH 7), 100 mM NaCl, 300 mM sucrose, 3 mM MgCl2, protease inhibitors), then rinsed twice quickly with CSK, and finally rinsed with PBS. Cells were immediately fixed in 2% paraformaldehyde in PBS for 20 min. Where indicated, after fixation, we performed a Click reaction according to the manufacturer’s instructions to reveal the EdU or EU (Click-iT EdU Alexa Fluor 488 imaging kit; Click-iT RNA Alexa Fluor 488 imaging kit, both from Invitrogen) to label nascent DNA or RNA respectively.

### Transfections and drug treatment

HeLa cells were transfected using lipofectamine RNAiMAX (Invitrogen) and small interfering RNAs (siRNA) were purchased from Dharmacon. We used ON-TARGETplus J-013610-06 (HIRA); ON-TARGETplus J-014195-05 (UBN1); ON-TARGETplus J-012454-09 (CABIN1); previously characterized siRNA (Groth *et al*. 2005; 2007) against ASF1a (GUGAAGAAUACGAUCAAGUUU) and ASF1b (CAACGAGUACCUC AACCCUUU). As siControl we used ON TARGETplus Non-targeting siRNA #1 (D-001810-01-05). For HIRA-YFP expressing plasmids, HeLa cells were transfected using Lipofectamine 2000 (Invitrogen). To arrest transcriptional activity, we used Flavopiridol hydrochloride hydrate (F3055) or Triptolide (T3652), both from Sigma-Aldrich, at 10uM concentration in complete medium. For long term treatment, to completely arrest transcription, cells were incubated with FLP during 3 hours prior to histone detection, and added in all steps to follow, for either old or new histone labelling pipeline.

### Antibodies

For immunofluorescence (IF), cells fixed on glass coversilps were blocked with 5% BSA in PBS-Tween 0.1% (PBST) for 45min at room temperature, and incubated with primary antibodies diluted in PBST-BSA for 45min. Primary antibodies were washed three times with PBST for 5min and cells were next incubated with secondary antibodies in PBST-BSA for 30min. Antibodies were washed three times in PBST as above, and cells were stained with DAPI. Antibodies were used at the following dilutions: anti-HIRA mouse monoclonal (WC119, Active Motif) IF 1:200 and Western Blot (WB) 1:100 (Hall *et al*. 2001); anti-RNAPIIS2ph (#04-1571, Millipore) WB and IF 1:500, anti-RNAPIIS5ph (#04-1572, Merck) WB and IF 1:500, anti-RNAPIIS7ph (61087, Active Motif) IF 1:300; anti-CABIN1 rabbit polyclonal (ab3349, Abcam) WB 1:1000; anti-UBN1 rabbit polyclonal (ab101282, Abcam) WB 1:1000; anti-HA epitope rat monoclonal (Roche) 1:1000; anti-ASF1a rabbit polyclonal (#2990, Cell Signaling) WB 1:1000; anti-ASF1b (Corpet *et al*. 2011); anti-DAXX rabbit monoclonal (D7810, Sigma) WB 1:4000; anti-p60 CAF-1 rabbit polyclonal WB 1:1000 (Green *et al*. 2010), anti-H3K36me3 (ab9050, Abcam) WB 1:1000; H4 mouse monoclonal (ab21830, Abcam) 1:1000.

### Epifluorescence microscopy and image analysis

For standard wide field epi-fluorescence imaging, coverslips were mounted in Vectashield medium. We used an AxioImager Zeiss Z1 microscope with a 63x or 100x objective.

#### Signal intensity quantification

FIJI (ImageJ) software was used to treat 2D images taken with the 63x objective and to quantify fluorescence signal within the nuclei area. To avoid misestimating histone loss due to cell cycle-related dilution effects, we quantified the fluorescence signal intensity normalized to the area of the nucleus as a proxy for DNA content. This process is automated using two FIJI macros, one for the subtraction of the background and another one for fluorescence quantification within the nuclei (Torné *et al*. 2018). Following quantification of EdU signal (Figure 1a) we systematically detected two clear populations of cells allowing to discriminate EdU positive/negative cells respectively in and outside of S-phase. Quantification of YFP following plasmid transfection for the rescue experiments (Figure 6), also allowed identification of a clear population of YFP positive cells, which were the only ones considering for TMR quantification. Quantification of the fluorescence intensity was performed in at least 100 nuclei per condition and in three independent experiments for all figures in the paper.

#### Spatial relationship analysis

For this analysis, 3D images were taken with the 100x objective and were deconvolved using Metamorph software. The blue channel (DAPI) is used to segment and separate the different nuclei. The spatial interactions between segmented foci (typically in the green channel) and the more homogeneous signal of (usually in the red channel) are estimated using a newly designed spatial statistic function inspired by (Helmuth *et al*. 2010; Lagache *et al*. 2013) and assessing the relationship between a point process and the spatial distribution of pixel intensities. The underlying concept is to observe the variations in intensity at an increasing distance *r* from foci. This function is normalized by the ratio between the local study volume and the nucleus total volume and is defined as:

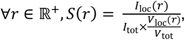

Where *r* is the study distance, *I*_loc_(*r*) and *I*_tot_ are respectively the local and total intensities, and *V*_loc_(*r*) and *V*_tot_ are respectively the local and total study volumes at a distance *r*, see Figure 2a) and 2b). At a high enough distance, the study volume is equal to the total nucleus volume, limiting edge effect. For *r* = 0, only the intensity of pixels under the segmented foci are accounted for.

Due to the volume normalization, a value *S*(*r*) = 1 indicates a total absence of interactions at scale *r* while values above show an increase in the intensity and values below 1 a depletion. In this study we are mostly interested in the trend of the function for small distances to see if the red signal is attracted or repulsed by the foci on the green channel. The function was first assessed on simulated data displaying perfect colocalization, spatial attraction and perfect anti-colocalization (see Figure 2c), and confidence intervals of departure from real independence between spots and intensities were defined by applying the function to positive and negative biological controls (Figure 2d), thus taking into account the potential biological confounding effects. The nucleoli were segmented and removed from the nuclei study volumes. The analysis is automated within a Fiji macro.

### Expression plasmids

The plasmids encoding HIRA-YFP WT and amino acid mutants were previously described in Ray-Gallet *et al.*, 2018.

### Cell extracts, immunoprecipitation and Western blotting

We prepared nuclear extract from HeLa cells as previously described (Martini *et al*. 1998), except that 300 mM NaCl was used. We obtained chromatin fraction by addition of benzonase to the pellet collected after nuclear extract preparation, followed by sonication. Immunoprecipitations were carried out overnight at 4°C with the appropriate primary antibody in the presence of 150 mM NaCl and 0.2% IGEPAL (Nonidet-P40 substitute) followed by an incubation with Dynabeads protein G (Invitrogen). For Western blot analysis, extracts or immunoprecipitated proteins were run on NuPAGE bis-tris 4-12% gels in MES or MOPS buffer (Invitrogen) and transferred to nitrocellulose membrane (Protran). Primary antibodies were detected using horse-radish-peroxidase conjugated secondary antibodies (Jackson ImmunoResearch Laboratories, Rockland for Trueblot) or protein A. We used SuperSignal West Pico or Dura chemiluminescent detection kits (Thermo Fischer) and the chemiluminescent signal was acquired using the ChemiDoc system equipped with an XRS camera (BioRad).

## Acknowledgements

We thank Cecilia Domrane for generating preliminary data with transcription inhibitors. We thank Jean-Pierre Quivy and Daniel Jeffery for critical reading of the manuscript, Patricia Le Baccon and the PICTIBiSA platform for help with the microscopy images acquisition and analysis. This work was supported by la Ligue Nationale contre le Cancer (Equipe labellisée Ligue), ANR-11-LABX-0044_DEEP and ANR-10-IDEX-0001-02 PSL, ANR-12-BSV5-0022-02 “CHAPINHIB”, ANR-14-CE16-0009 “Epicure”, ANR-14-CE10-0013 “CELLECTCHIP”, EU project 678563 “EPOCH28”, ERC-2015-ADG-694694 “ChromADICT”, ANR-16-CE15-0018 “CHRODYT”, ANR-16-CE12-0024 “CHIFT”, ANR-16-CE11-0028 “REPLICAF”, PSL-AFdL “TRACK”’.

## Author contributions

G.A. and G.A.O. supervised the work. G.A., G.A.O. and J.T. conceived the strategy and wrote the paper. J.T. performed epifluorescence and cell biology experiments. J.T and G.A.O analysed the data. D.R.G. designed and performed the biochemistry experiments. E.B. carried out transcription experiments with inhibitors drugs. A.C. performed fits and interpretation of kinetic data. M.G. designed imaging analysis methods. Critical reading and discussion of all data involved all authors.

---

**Supplementary Figure 1.**
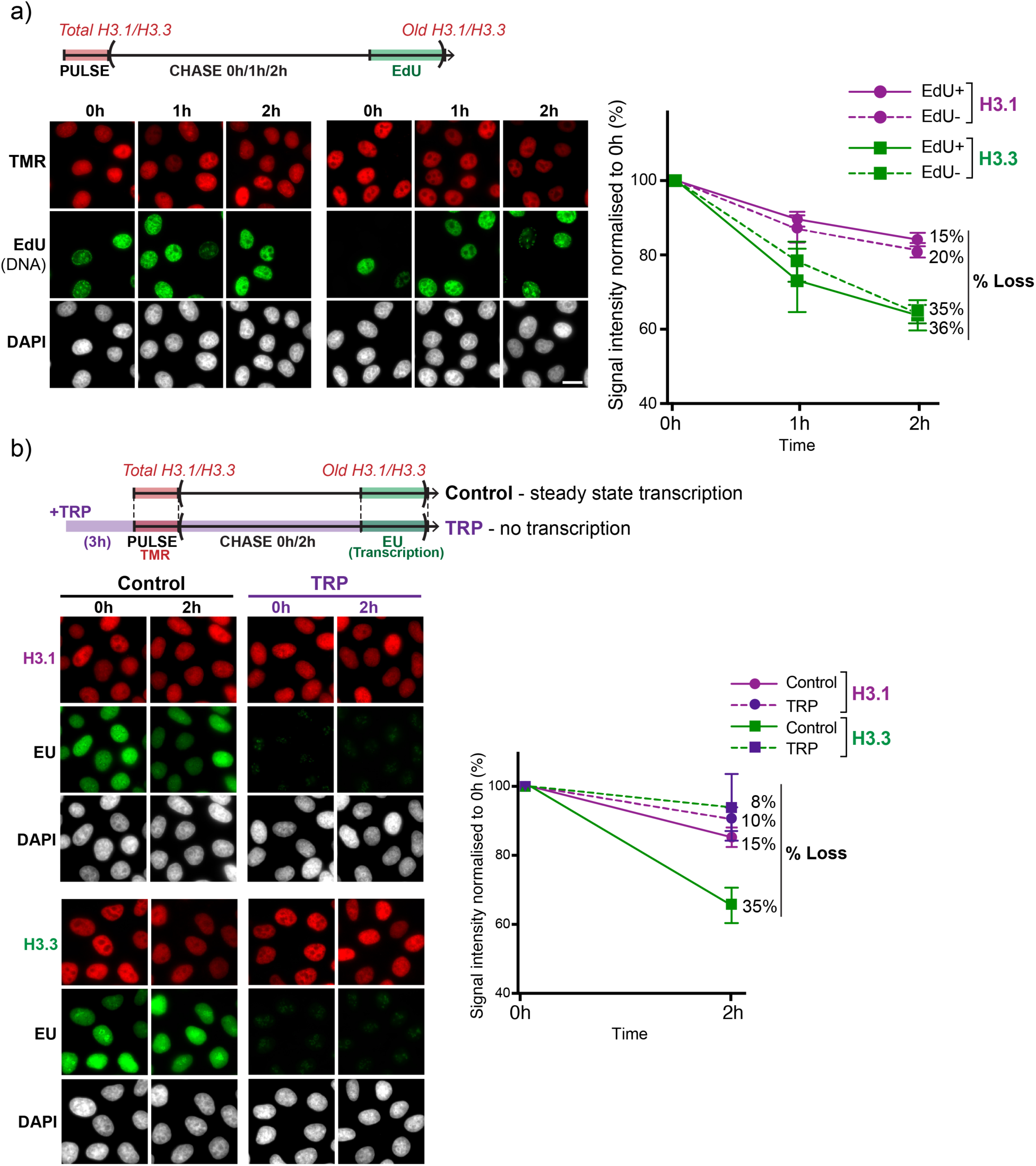
Transcription inhibited by Triptolide also prevents H3.3 loss. **a)** Top: Experimental set-up to track total or old H3.1/H3.3 using SNAP-tag labelling. A pulse using the fluorophore TMR (red) labels SNAP-tagged H3.3 or H3.1, cells are triton-extracted and fixed at different chase times to reveal total (0h) or old (1h, 2h) chromatin-bound histones. EdU labelling (green) marks nascent DNA allowing identification of cells in or outside S-phase, while DNA is stained with DAPI (grey). Bottom-left: representative wide field epifluorescence images of H3.1- or H3.3-SNAP after 0h, 1h and 2h of chase time. Bottom-right: quantification of average nuclear TMR signal for H3.1 (purple) and H3.3 (green), expressed as a percentage of the average value at chase time 0h. The average percentage of loss at 2h chase time is indicated for each sample. Cells were grouped as EdU positive (EdU+: cells in S-phase, full lines) or EdU negative (EdU-: cells outside of S-phase, dashed lines). **b)** Top: Experimental set-up to track total and old H3.1/H3.3-SNAP (TMR, red) in the presence or absence of the transcription inhibitor Triptolide (TRP), performed in parallel to Flavopiridol treatment in Figure 1b. EU labelling (green) marks nascent RNA and is used to confirm the absence of transcription in Triptolide-treated cells. Bottom-left: representative wide field epifluorescence images of control and TRP-treated H3.3- or H3.1-SNAP cells after 0h and 2h of TMR chase time. Bottom-right: quantification of average TMR signal for H3.3-(green) and H3.1-SNAP (purple) of untreated (full lines) or TRP-treated (dashed lines) cells, as a percentage of the average value at time 0h for each condition. For all samples, n>200 nuclei. Plots show average and standard error for three biological replicates. Control data is the same as in Figure 1b. Scale bars represent 10 μm.

**Supplementary Figure 2.**
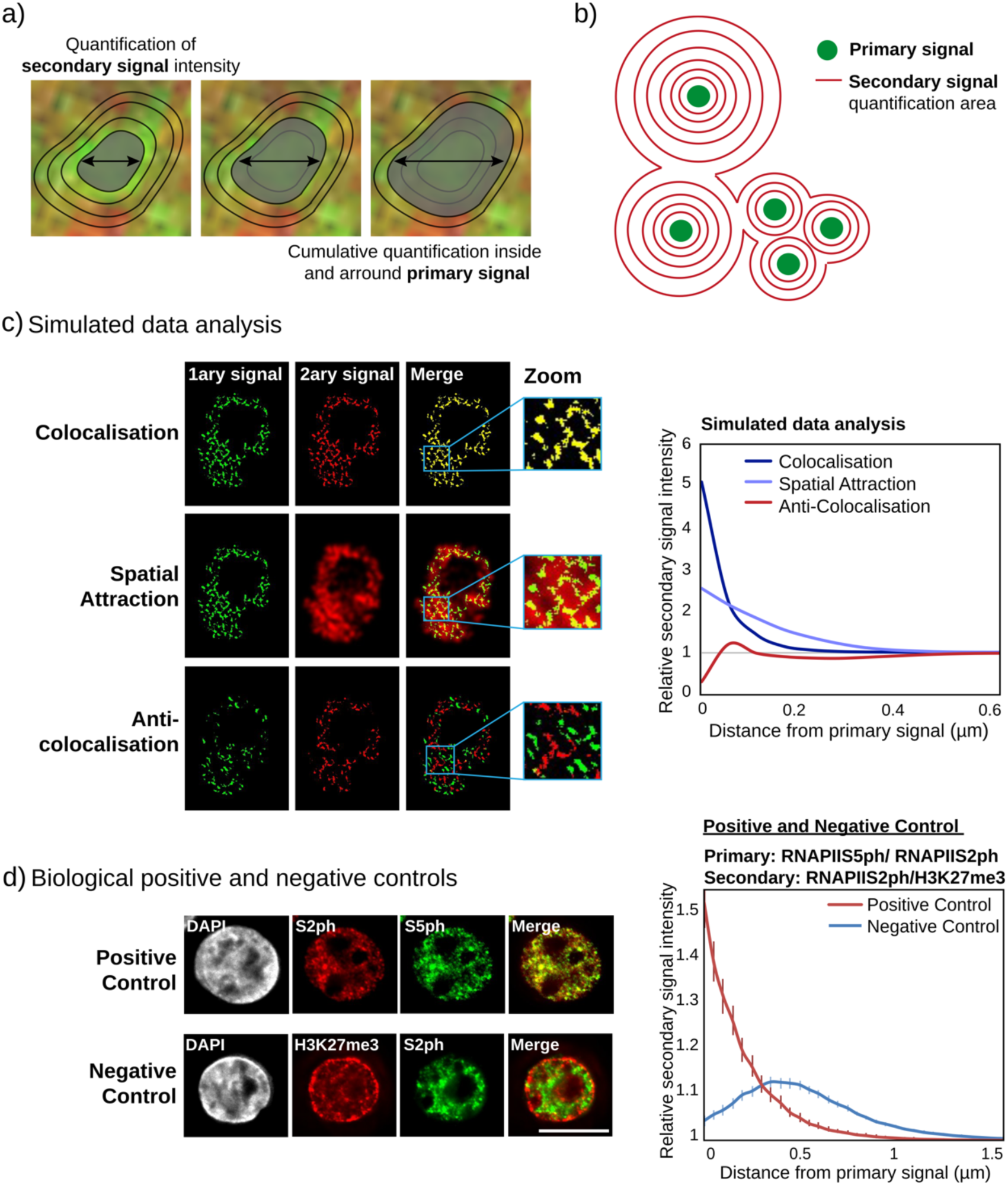
Image analysis methodology for spatial relationship. **a)** Scheme showing the area considered for cumulative secondary signal quantification inside and around primary signal foci. **b)** Scheme showing how secondary signal is assigned to a single primary signal spot. **c)** Simulated data images and corresponding analysis plot, for cases of perfect colocalization, spatial attraction and perfect anticolocalisation cases. **d)** Left top: representative deconvolved epifluorescence images of cells stained for RNAPIIS2ph (red) and RNAPIIS5ph (green) as positive biological control for colocalization. Left bottom: representative deconvolved epifluorescence images of cells stained for RNAPIIS2ph (green) and H3K27me3 (red) as negative biological control for colocalization. DNA is stained with DAPI (grey). Right: spatial relationship analysis showing colocalization of RNAPIIS2ph and RNAPIIS5ph and anticolocalisation for RNAPIIS2ph and H3K27me3. For spatial relationship analysis, numbers are averages from n>40 cells. Scale bar represents 10µm.

**Supplementary Figure 3.**
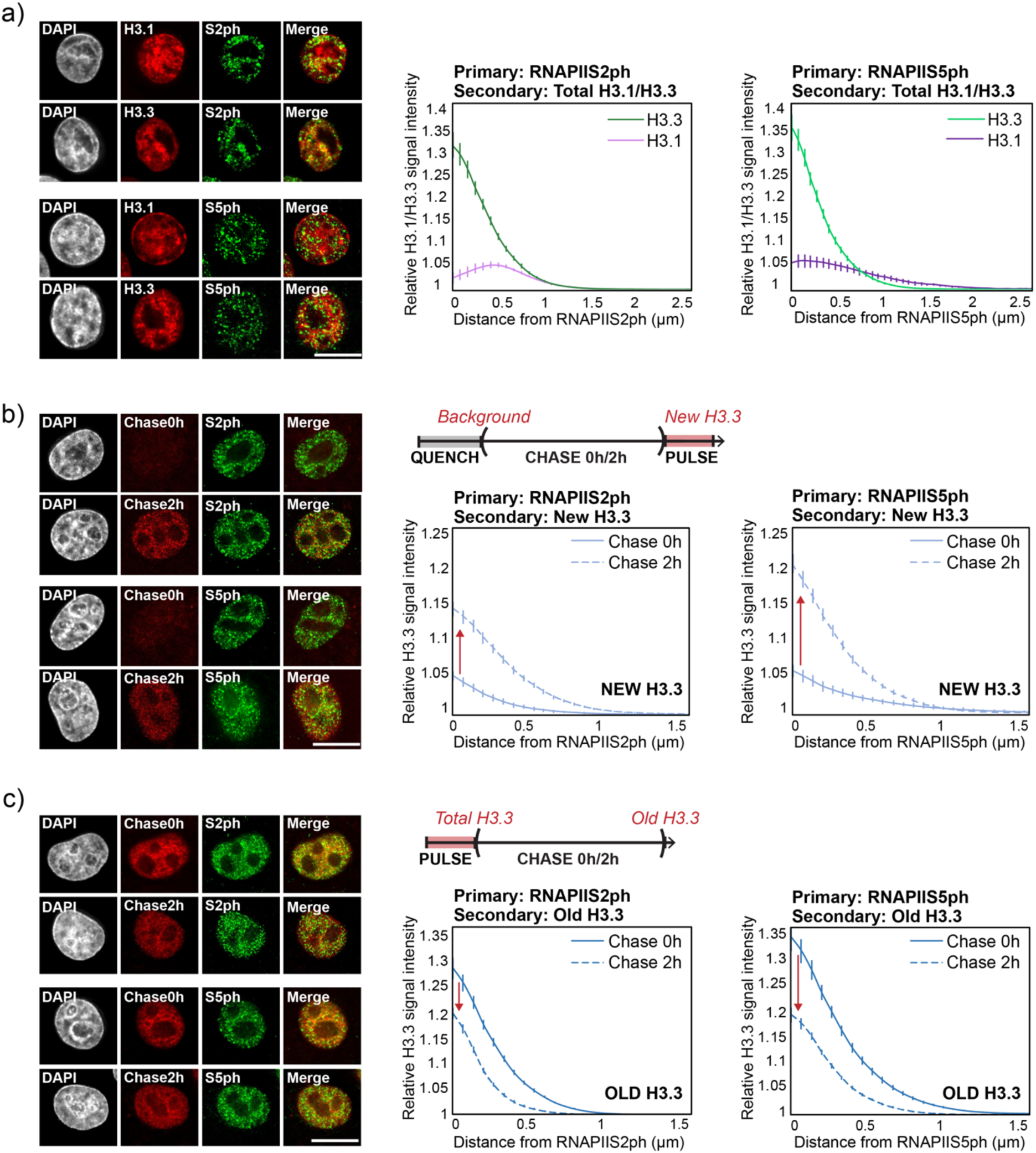
H3.3 is dynamically exchanged at nuclear domains marked by RNAPIIS2ph and RNAPIIS5ph. **a)** Left: representative deconvolved epifluorescence images of cells stained for global H3.1- or H3.3-SNAP (TMR, red), RNAPIIS2ph or RNAPIIS5ph (green) and DNA (DAPI, grey). Right: spatial relationship analysis showing an enrichment of H3.3 (green) and depletion of H3.1 (purple) at RNAPIIS2ph and RNAPIIS5ph foci. **b)** Top right: Experimental set-up to track new H3.3-SNAP histones synthesized after 0h and 2h of chase time. Left: representative deconvolved epiflorescence images of cells stained for new H3.1- or H3.3-SNAP (TMR, red), RNAPIIS2ph or RNAPIIS5ph (green) and DNA (DAPI, white). Bottom right: spatial relationship analysis of new H3.3 distribution relative to RNAPIIS2ph and RNAPIIS5ph at 0h (full line) and 2h (dashed line) chase time showing accumulation of new histone preferentially at these foci. **c)** as in b), except old H3.3-SNAP was tracked, showing loss of old histone preferentially at these foci. All plots show average and standard error of n>40 cells from one representative biological replicate. Scale bars represent 10 μm.

**Supplementary Figure 4.**
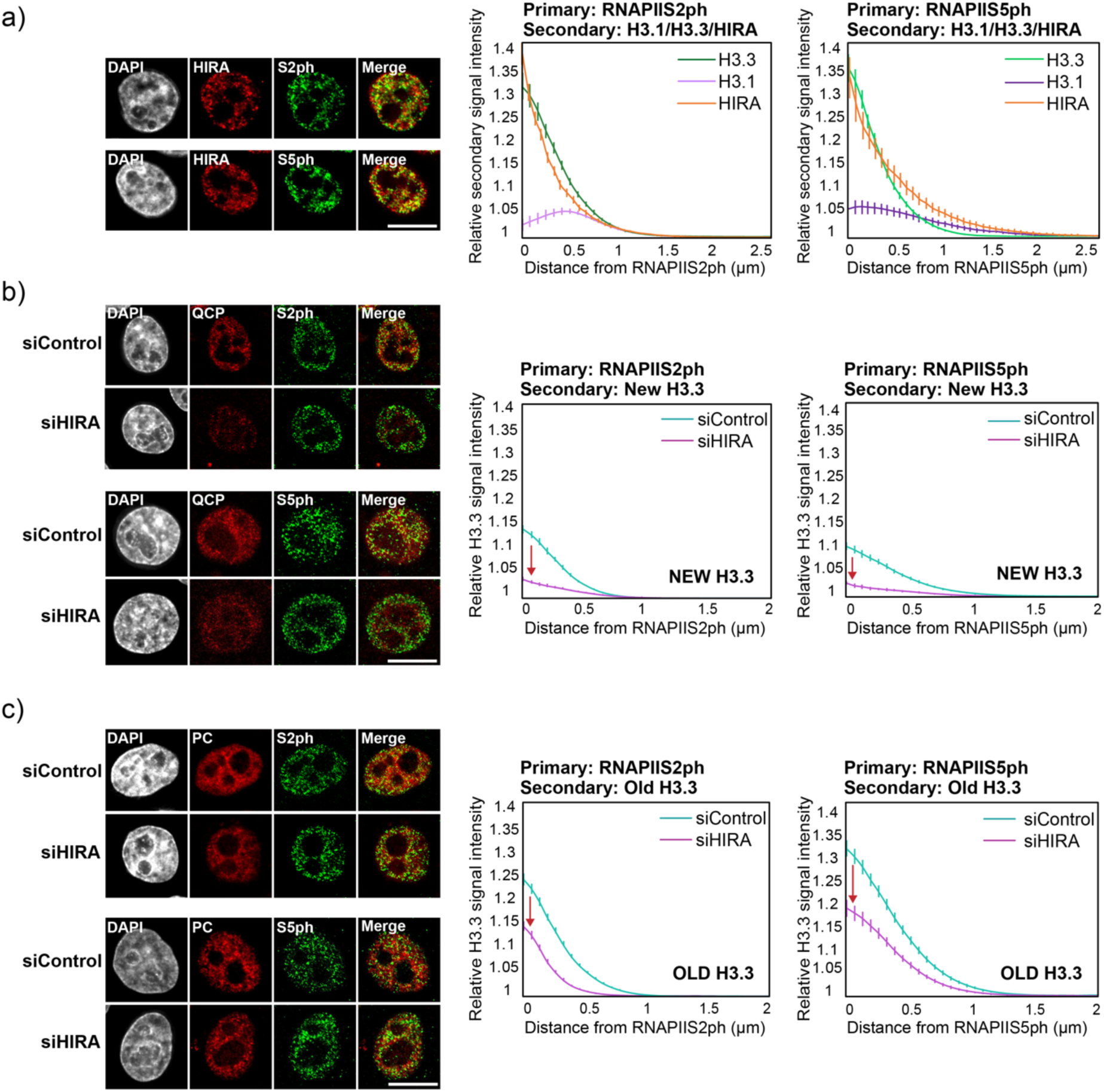
HIRA controls both deposition of new H3.3 and recycling of old H3.3 at RNAPIIS2ph and RNAPIIS5ph foci. **a)** Left: representative deconvolved epifluorescence images for cells stained for total HIRA (TMR, red) together with RNAPIIS2ph (green, top) or RNAPIIS5ph (green, bottom) and DNA (DAPI, grey). Right: Spatial relationship analysis showing enrichment of total H3.3 (green) and HIRA (orange), and depletion of total H3.1 (purple), at RNAPIIS2ph or RNAPIIS5ph foci. **b)** Left: representative deconvolved epifluorescence images for cells stained for new H3.3-SNAP (TMR, red) after 72h siControl or siHIRA together with RNAPIIS2ph (green, top) or RNAPIIS5ph (green, bottom) and DNA (DAPI, grey). Right: spatial relationship analysis showing depletion of new H3.3 at RNAPIIS2ph or RNAPIIS5 foci upon HIRA knockdown (purple lines) compared to control (blue lines). **c)** as in b), except old H3.3-SNAP was tracked, showing depletion of old H3.3 at RNAPIIS2ph and RNAPIIS5ph foci in HIRA knockdown cells. All plots show average and standard error of n>40 cells from one representative biological replicate. Scale bars represent 10 μm.

**Supplementary Figure 5.**
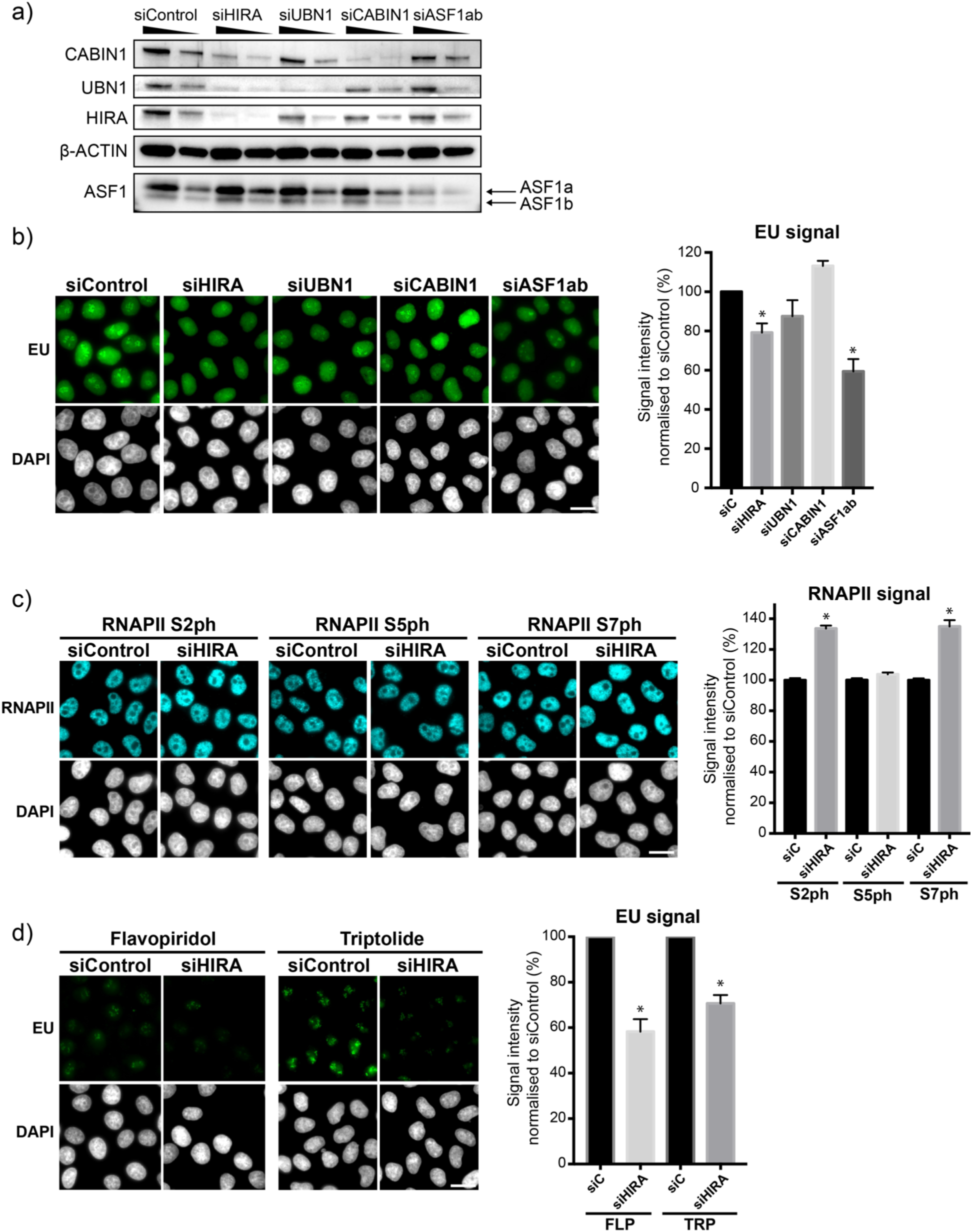
Effects of HIRA Knockdown on transcription efficiency. **a)** Western blot of total protein extracts showing efficient 72h siRNA knockdown treatment targeting different chaperones Note that, as previously described, HIRA knockdown entails depletion of HIRA, UBN1 and CABIN1. **b)** HIRA, UBN1 and ASF1 knockdown leads to decrease in nascent transcription. Left: representative epifluorescence images of cells stained for EU (nascent RNA, green) and DAPI (grey) following 72h knockdown using siRNA targeting HIRA, UBN1, CABIN1 or ASF1ab or a control. Right: quantification of average EU signal (green), normalized to average signal in siControl, showing decreased nascent transcription upon HIRA, UBN1 or ASF1 knockdown. **c)** Global increase in RNAPIIS2ph and RNAPIIS7ph, but not RNAPIIS5ph upon HIRA knockdown. Left: representative epifluorescence images of cells stained for RNAPIIS2ph, RNAPIIS5ph and RNAPIIS7ph (cyan) and DAPI (grey) after siControl or siHIRA treatment. Left: quantification of average RNAPII signal, normalized to average signal in siControl, revealing an increase in S2ph and S7ph RNAPII forms upon HIRA knockdown. **d)** Flavopiridol and Triptolide are even more efficient upon HIRA knockdown. Left: representative epifluorescence images of cells stained for EU (green) and DAPI (grey) following 3h or Flavopiridol (FLP) or Triptolide (TRP) treatment and after 72h siControl or siHIRA knockdown treatment. Right: quantification of average EU signal (green), normalized to siControl, showing decreased signal upon HIRA knockdown. All plots show average and standard error for n>200 cells from two biological replicates. Standard t-test showed statistical significance (*p < 0.05).

**Supplementary Figure 6.**
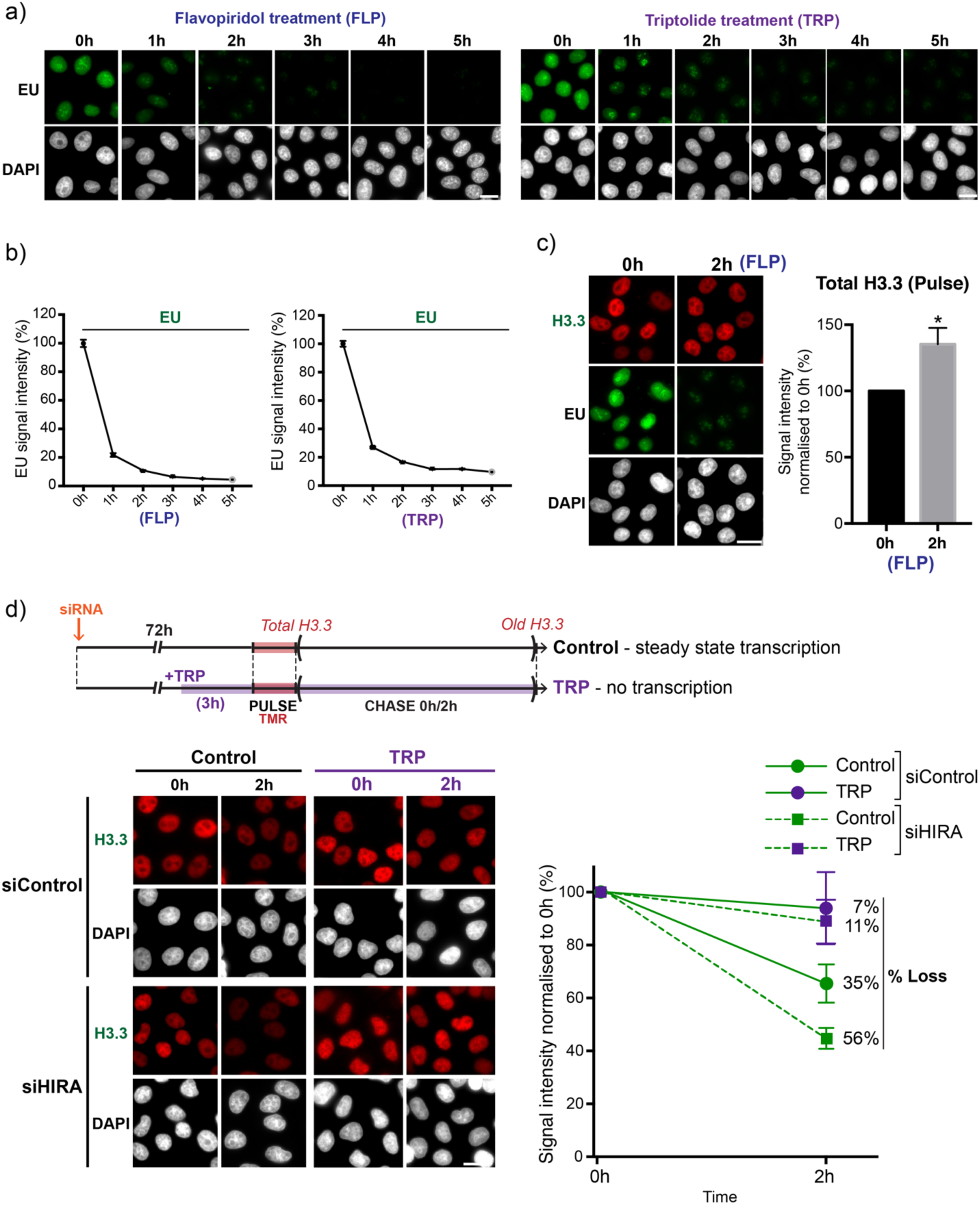
Effects of RNAPII-inhibiting drugs on H3.3. **a)** Representative images of cells treated with Flavopiridol (FLP) or Triptolide (TRP) for the indicated times and labelled with EU (green) to measure nascent transcript levels and DAPI (DNA, grey). **b)** Corresponding quantifications of total nuclear EU signal normalized to untreated cells (0h) allowed to distinguish between full signal in steady state transcription, to no signal after complete transcription shutdown (3h-5h). **b)** Left: representative epifluorescence images of cells stained for total H3.3-SNAP (TMR, red) EU (nascent RNA, green) and DAPI (grey) after 0h or 2h of FLP treatment. Right: quantification of average TMR signal (red), as a percentage of average values at 0h, showing an increase in total H3.3 upon transcriptional arrest. **b)** Same as in Figure 4, excerpt Triptolide (TRP) was used to inhibit transcriptiopn. Left: representative images of total (0h) or old (2h) H3.3-SNAP (TMR, red), in control conditions or during TRP treatment and 72h following knockdown using Control or HIRA-targeting siRNA. Right: quantification of TMR signal for control (full lines) and HIRA knockdown (dashed lines), untreated (green) and TRP-treated (purple) cells. As for FLP treatment, the results indicate that transcriptional activity is necessary to reveal the effect of HIRA knockdown on H3.3 recycling. Control data is the same as in Figure 4. All plots show average and standard error for n>200 cells from two biological replicates. Standard t-test indicated statistical significance (*: p < 0.05). Scale bars represent 10 μm.

**Supplementary Figure 7.**
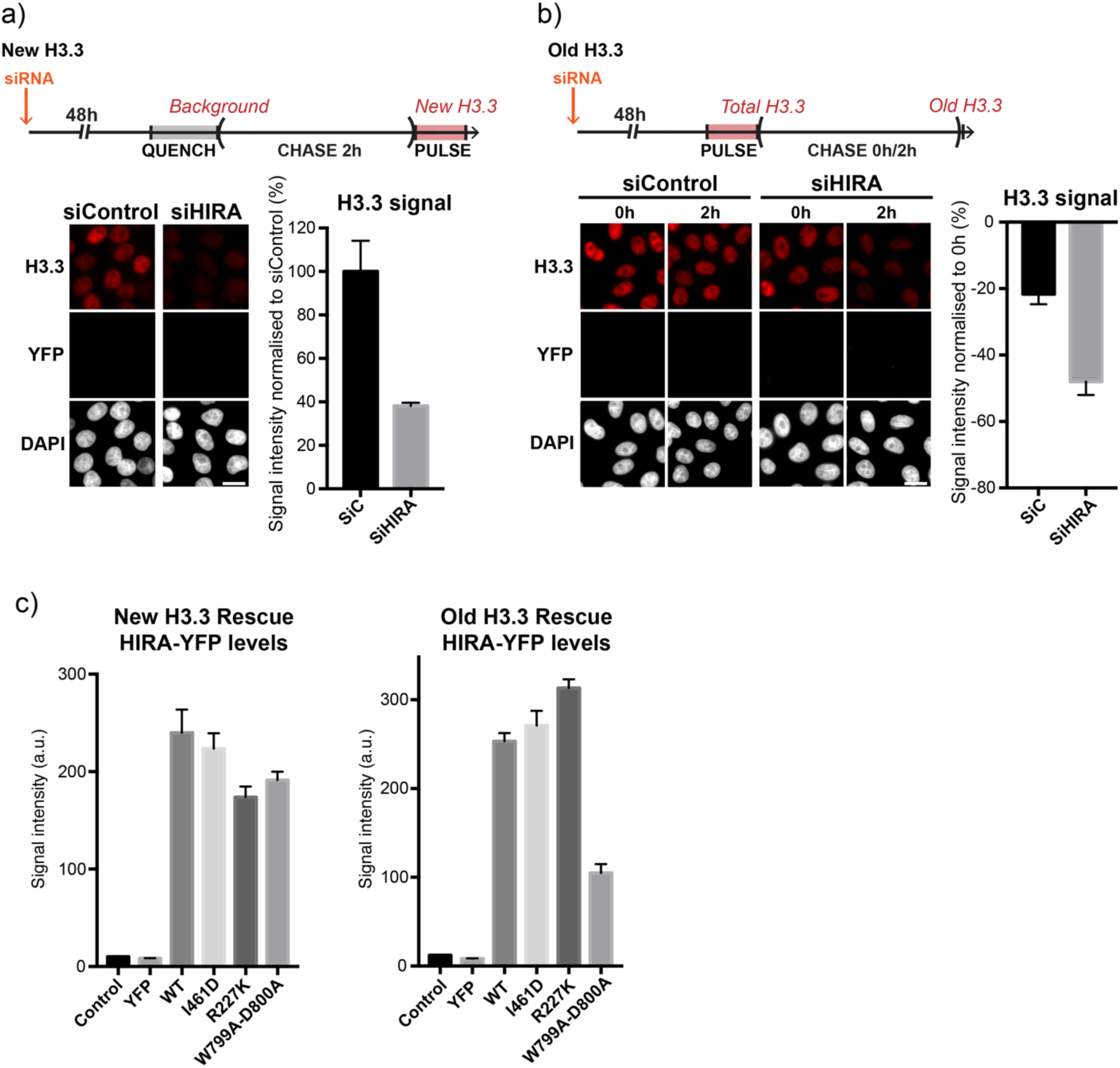
Expression of HIRA-YFP mutant transgenic constructs. **a)** Top: Experimental set-up to track new H3.3-SNAP (TMR, red) under siControl or SiHIRA. Bottom-left: representative wide field epifluorescence images of new H3.3-SNAP (red) and DAPI (grey) following 72h siRNA Control or HIRA. Bottom-right: quantification of average TMR signal for H3.3-SNAP (red) normalized to siControl. As expected, a YFP transgenic protein is not detected in the nucleus after permeabilization: this is used as a negative control for HIRA-YFP rescue experiments in Figure 6. For all samples, n>200 nuclei. **b)** as in a) but for old H3.3-SNAP. Plot shows 2h old H3.3-SNAP quantification under siControl or siHIRA normalized to 0h. Plots show average and standard error for two biological replicates. Scale bars represent 10 μm. **c)** HIRA-YFP wild type (WT) and mutants are present at comparable levels in the nucleus following transfection. Fluorescence quantification of YFP in HIRA knockdown cells expressing HIRA-YFP WT and HIRA-YFP mutants following 48h of siHIRA treatment in rescue experiments for new H3.3 deposition (left) and the old H3.3 loss (right). Plots show average and standard error for n>200 cells from a single experiment.

## REFERENCES

Adam, Salomé, Sophie E. Polo, and Geneviève Almouzni. 2013. “Transcription Recovery after DNA Damage Requires Chromatin Priming by the H3.3 Histone Chaperone HIRA.” Cell 155 (1): 94–106. https://doi.org/10.1016/j.cell.2013.08.029.

Ahmad, Kami, and Steven Henikoff. 2002. “The Histone Variant H3.3 Marks Active Chromatin by Replication-Independent Nucleosome Assembly.” Molecular Cell 9 (6): 1191–1200. https://doi.org/10.1016/S1097-2765(02)00542-7.

Baldi, Sandro, Stefan Krebs, and Peter B. Becker. 2018. “Genome-Wide Measurement of Local Nucleosome Array Regularity and Spacing by Nanopore Sequencing.” Molecular Biology 25: 11.

Bannister, Andrew J., Robert Schneider, Fiona A. Myers, Alan W. Thorne, Colyn Crane-Robinson, and Tony Kouzarides. 2005. “Spatial Distribution of Di- and Tri-Methyl Lysine 36 of Histone H3 at Active Genes.” Journal of Biological Chemistry 280 (18): 17732–36. https://doi.org/10.1074/jbc.M500796200.

Banumathy, G., N. Somaiah, R. Zhang, Y. Tang, J. Hoffmann, M. Andrake, H. Ceulemans, D. Schultz, R. Marmorstein, and P. D. Adams. 2009. “Human UBN1 Is an Ortholog of Yeast Hpc2p and Has an Essential Role in the HIRA/ASF1a Chromatin-Remodeling Pathway in Senescent Cells.” Molecular and Cellular Biology 29 (3): 758–70. https://doi.org/10.1128/MCB.01047-08.

Boehning, Marc, Claire Dugast-Darzacq, Marija Rankovic, Anders S. Hansen, Taekyung Yu, Herve Marie-Nelly, David T. McSwiggen, et al. 2018. “RNA Polymerase II Clustering through Carboxy-Terminal Domain Phase Separation.” Nature Structural & Molecular Biology 25 (9): 833–40. https://doi.org/10.1038/s41594-018-0112-y.

Bondarenko, Vladimir A., Louise M. Steele, Andrea Újvári, Daria A. Gaykalova, Olga I. Kulaeva, Yury S. Polikanov, Donal S. Luse, and Vasily M. Studitsky. 2006. “Nucleosomes Can Form a Polar Barrier to Transcript Elongation by RNA Polymerase II.” Molecular Cell 24 (3): 469–79. https://doi.org/10.1016/j.molcel.2006.09.009.

Chen, Shoudeng, Anne Rufiange, Hongda Huang, Kanagalaghatta R. Rajashankar, Amine Nourani, and Dinshaw J. Patel. 2015. “Structure–Function Studies of Histone H3/H4 Tetramer Maintenance during Transcription by Chaperone Spt2.” Genes & Development 29 (12): 1326–40. https://doi.org/10.1101/gad.261115.115.

Cho, Won-Ki, Jan-Hendrik Spille, Micca Hecht, Choongman Lee, Charles Li, Valentin Grube, and Ibrahim I. Cisse. 2018. “Mediator and RNA Polymerase II Clusters Associate in Transcription-Dependent Condensates.” Science 361 (6400): 412–15. https://doi.org/10.1126/science.aar4199.

Clément, Camille, Guillermo A. Orsi, Alberto Gatto, Ekaterina Boyarchuk, Audrey Forest, Bassam Hajj, Judith Miné-Hattab, et al. 2018. “High-Resolution Visualization of H3 Variants during Replication Reveals Their Controlled Recycling.” Nature Communications 9 (1): 3181. https://doi.org/10.1038/s41467-018-05697-1.

Cook, Adam J.L., Zachary A. Gurard-Levin, Isabelle Vassias, and Geneviève Almouzni. 2011. “A Specific Function for the Histone Chaperone NASP to Fine-Tune a Reservoir of Soluble H3-H4 in the Histone Supply Chain.” Molecular Cell 44 (6): 918–27. https://doi.org/10.1016/j.molcel.2011.11.021.

Corpet, Armelle, Leanne De Koning, Joern Toedling, Alexia Savignoni, Frédérique Berger, Charlène Lemaître, Roderick J O’Sullivan, et al. 2011. “Asf1b, the Necessary Asf1 Isoform for Proliferation, Is Predictive of Outcome in Breast Cancer: Specific Importance of Asf1b in Proliferation.” The EMBO Journal 30 (3): 480–93. https://doi.org/10.1038/emboj.2010.335.

Daganzo, Sally M, Jan P Erzberger, Wendy M Lam, Emmanuel Skordalakes, Rugang Zhang, Alexa A Franco, Steven J Brill, Peter D Adams, James M Berger, and Paul D Kaufman. 2003. “Structure and Function of the Conserved Core of Histone Deposition Protein Asf1.” Current Biology 13 (24): 2148–58. https://doi.org/10.1016/j.cub.2003.11.027.

Deal, R. B., J. G. Henikoff, and S. Henikoff. 2010. “Genome-Wide Kinetics of Nucleosome Turnover Determined by Metabolic Labeling of Histones.” Science 328 (5982): 1161–64. https://doi.org/10.1126/science.1186777.

Deaton, Aimee M, Mariluz Gómez-Rodríguez, Jakub Mieczkowski, Michael Y Tolstorukov, Sharmistha Kundu, Ruslan I Sadreyev, Lars ET Jansen, and Robert E Kingston. 2016. “Enhancer Regions Show High Histone H3.3 Turnover That Changes during Differentiation.” ELife 5 (June): e15316. https://doi.org/10.7554/eLife.15316.

Dion, M. F., T. Kaplan, M. Kim, S. Buratowski, N. Friedman, and O. J. Rando. 2007. “Dynamics of Replication-Independent Histone Turnover in Budding Yeast.” Science 315 (5817): 1405–8. https://doi.org/10.1126/science.1134053.

Drane, P., K. Ouararhni, A. Depaux, M. Shuaib, and A. Hamiche. 2010. “The Death-Associated Protein DAXX Is a Novel Histone Chaperone Involved in the Replication-Independent Deposition of H3.3.” Genes & Development 24 (12): 1253–65. https://doi.org/10.1101/gad.566910.

Edmunds, John, Mahadevan, and mahadevan. 2008. “Dynamic Histone H3 Methylation during Gene Induction: HYPB/Setd2 Mediates All H3K36 Trimethylation.” EMBO J 27(2): 406–420. https://doi.org/10.1038/sj.emboj.7601967

English, Christine M., Melissa W. Adkins, Joshua J. Carson, Mair E.A. Churchill, and Jessica K. Tyler. 2006. “Structural Basis for the Histone Chaperone Activity of Asf1.” Cell 127 (3): 495–508. https://doi.org/10.1016/j.cell.2006.08.047.

Farnung, Lucas, Seychelle M. Vos, and Patrick Cramer. 2018. “Structure of Transcribing RNA Polymerase II-Nucleosome Complex.” Nature Communications 9 (1): 5432. https://doi.org/10.1038/s41467-018-07870-y.

Gaillard, Pierre-Henri L, Emmanuelle M.-D Martini, Paul D Kaufman, Bruce Stillman, Ethel Moustacchi, and Geneviève Almouzni. 1996. “Chromatin Assembly Coupled to DNA Repair: A New Role for Chromatin Assembly Factor I.” Cell 86 (6): 887–96. https://doi.org/10.1016/S0092-8674(00)80164-6.

Gan, Haiyun, Albert Serra-Cardona, Xu Hua, Hui Zhou, Karim Labib, Chuanhe Yu, and Zhiguo Zhang. 2018. “The Mcm2-Ctf4-Polα Axis Facilitates Parental Histone H3-H4 Transfer to Lagging Strands.” Molecular Cell 72 (1): 140–151.e3. https://doi.org/10.1016/j.molcel.2018.09.001.

Ghamari, A., M. P. C. van de Corput, S. Thongjuea, W. A. van Cappellen, W. van IJcken, J. van Haren, E. Soler, D. Eick, B. Lenhard, and F. G. Grosveld. 2013. “In Vivo Live Imaging of RNA Polymerase II Transcription Factories in Primary Cells.” Genes & Development 27 (7): 767–77. https://doi.org/10.1101/gad.216200.113.

Goldberg, Aaron D., Laura A. Banaszynski, Kyung-Min Noh, Peter W. Lewis, Simon J. Elsaesser, Sonja Stadler, Scott Dewell, et al. 2010. “Distinct Factors Control Histone Variant H3.3 Localization at Specific Genomic Regions.” Cell 140 (5): 678–91. https://doi.org/10.1016/j.cell.2010.01.003.

Green, Erin M, Andrew J Antczak, Aaron O Bailey, Alexa A Franco, Kevin J Wu, R Yates, and Paul D Kaufman. 2010. “Replication-Independent Histone Deposition by the HIR Complex and Asf,” Current Biology 15(22):2044–9. https://doi.org/10.1016/j.cub.2005.10.053

Gregersen, Lea H., and Jesper Q. Svejstrup. 2018. “The Cellular Response to Transcription-Blocking DNA Damage.” Trends in Biochemical Sciences 43 (5): 327–41. https://doi.org/10.1016/j.tibs.2018.02.010.

Groth, Anja, Dominique Ray-Gallet, Jean-Pierre Quivy, Jiri Lukas, Jiri Bartek, and Geneviève Almouzni. 2005. “Human Asf1 Regulates the Flow of S Phase Histones during Replicational Stress.” Molecular Cell 17 (2): 301–11. https://doi.org/10.1016/j.molcel.2004.12.018.

Groth, Anja, Walter Rocha, Alain Verreault, and Geneviève Almouzni. 2007. “Chromatin Challenges during DNA Replication and Repair.” Cell 128 (4): 721–33. https://doi.org/10.1016/j.cell.2007.01.030.

Guan, Ling, Peng He, Fang Yang, Yuan Zhang, Yunfei Hu, Jienv Ding, Yu Hua, et al. 2017. “Sap1 Is a Replication-Initiation Factor Essential for the Assembly of Pre-Replicative Complex in the Fission Yeast *Schizosaccharomyces Pombe*.” Journal of Biological Chemistry 292 (15): 6056–75. https://doi.org/10.1074/jbc.M116.767806.

Guo, Yang Eric, John C. Manteiga, Jonathan E. Henninger, Benjamin R. Sabari, Alessandra Dall’Agnese, Nancy M. Hannett, Jan-Hendrik Spille, et al. 2019. “Pol II Phosphorylation Regulates a Switch between Transcriptional and Splicing Condensates.” Nature 572 (7770): 543–48. https://doi.org/10.1038/s41586-019-1464-0.

Gurard-Levin, Zachary A., Jean-Pierre Quivy, and Geneviève Almouzni. 2014. “Histone Chaperones: Assisting Histone Traffic and Nucleosome Dynamics.” Annual Review of Biochemistry 83 (1): 487–517. https://doi.org/10.1146/annurev-biochem-060713-035536.

Hall, C., D. M. Nelson, X. Ye, K. Baker, J. A. DeCaprio, S. Seeholzer, M. Lipinski, and P. D. Adams. 2001. “HIRA, the Human Homologue of Yeast Hir1p and Hir2p, Is a Novel Cyclin-Cdk2 Substrate Whose Expression Blocks S-Phase Progression.” Molecular and Cellular Biology 21 (5): 1854–65. https://doi.org/10.1128/MCB.21.5.1854-1865.2001.

Helmuth, Jo A, Grégory Paul, and Ivo F Sbalzarini. 2010. “Beyond Co-Localization: Inferring Spatial Interactions between Sub-Cellular Structures from Microscopy Images.” BMC Bioinformatics 11 (1): 372. https://doi.org/10.1186/1471-2105-11-372.

Horard, Béatrice, Laure Sapey-Triomphe, Emilie Bonnefoy, and Benjamin Loppin. 2018. “ASF1 Is Required to Load Histones on the HIRA Complex in Preparation of Paternal Chromatin Assembly at Fertilization.” Epigenetics & Chromatin 11 (1): 19. https://doi.org/10.1186/s13072-018-0189-x.

Janicki, Susan M, Toshiro Tsukamoto, Simone E Salghetti, William P Tansey, Ravi Sachidanandam, Kannanganattu V Prasanth, Thomas Ried, et al. 2004. “From Silencing to Gene Expression: Real-Time Analysis in Single Cells,” 16.

Jeronimo, Célia, Christian Poitras, and François Robert. 2019. “Histone Recycling by FACT and Spt6 during Transcription Prevents the Scrambling of Histone Modifications.” Cell Reports 28 (5): 1206–1218.e8. https://doi.org/10.1016/j.celrep.2019.06.097.

Keppler, Antje, Susanne Gendreizig, Thomas Gronemeyer, Horst Pick, Horst Vogel, and Kai Johnsson. 2003. “A General Method for the Covalent Labeling of Fusion Proteins with Small Molecules in Vivo.” Nature Biotechnology 21 (1): 86–89. https://doi.org/10.1038/nbt765.

Kireeva, Maria L., Brynne Hancock, Gina H. Cremona, Wendy Walter, Vasily M. Studitsky, and Mikhail Kashlev. 2005. “Nature of the Nucleosomal Barrier to RNA Polymerase II.” Molecular Cell 18 (1): 97–108. https://doi.org/10.1016/j.molcel.2005.02.027.

Kireeva, Maria L, Wendy Walter, Vladimir Tchernajenko, Vladimir Bondarenko, Mikhail Kashlev, and Vasily M Studitsky. 2002. “Nucleosome Remodeling Induced by RNA Polymerase II.” Molecular Cell 9 (3): 541–52. https://doi.org/10.1016/S1097-2765(02)00472-0.

Knezetic, Joseph A., and Donal S. Luse. 1986. “The Presence of Nucleosomes on a DNA Template Prevents Initiation by RNA Polymerase II in Vitro.” Cell 45 (1): 95–104. https://doi.org/10.1016/0092-8674(86)90541-6.

Kraushaar, Daniel C, Wenfei Jin, Alika Maunakea, Brian Abraham, Misook Ha, and Keji Zhao. 2013. “Genome-Wide Incorporation Dynamics Reveal Distinct Categories of Turnover for the Histone Variant H3.3.” Genome Biology 14 (10): R121. https://doi.org/10.1186/gb-2013-14-10-r121.

Kujirai, Tomoya, Haruhiko Ehara, Yuka Fujino, Mikako Shirouzu, Shun-ichi Sekine, and Hitoshi Kurumizaka. 2018. “Structural Basis of the Nucleosome Transition during RNA Polymerase II Passage.” Science 362 (6414): 595–98. https://doi.org/10.1126/science.aau9904.

Kujirai, Tomoya, and Hitoshi Kurumizaka. 2020. “Transcription through the Nucleosome.” Current Opinion in Structural Biology 61 (April): 42–49. https://doi.org/10.1016/j.sbi.2019.10.007.

Kulaeva, O. I., F.-K. Hsieh, and V. M. Studitsky. 2010. “RNA Polymerase Complexes Cooperate to Relieve the Nucleosomal Barrier and Evict Histones.” Proceedings of the National Academy of Sciences 107 (25): 11325–30. https://doi.org/10.1073/pnas.1001148107.

Kulaeva, Olga I, Daria A Gaykalova, Nikolai A Pestov, Viktor V Golovastov, Dmitry G Vassylyev, Irina Artsimovitch, and Vasily M Studitsky. 2009. “Mechanism of Chromatin Remodeling and Recovery during Passage of RNA Polymerase II.” Nature Structural & Molecular Biology 16 (12): 1272–78. https://doi.org/10.1038/nsmb.1689.

Lagache, Thibault, Gabriel Lang, Nathalie Sauvonnet, and Jean-Christophe Olivo-Marin. 2013. “Analysis of the Spatial Organization of Molecules with Robust Statistics.” Edited by Joshua Z. Rappoport. PLoS ONE 8 (12): e80914. https://doi.org/10.1371/journal.pone.0080914.

Lee, Cheol-Koo, Yoichiro Shibata, Bhargavi Rao, Brian D Strahl, and Jason D Lieb. 2004. “Evidence for Nucleosome Depletion at Active Regulatory Regions Genome-Wide.” Nature Genetics 36 (8): 900–905. https://doi.org/10.1038/ng1400.

Lewis, Peter W, Simon J Elsaesser, Kyung-Min Noh, Sonja C Stadler, and C David Allis. 2010. “Daxx Is an H3.3-Specific Histone Chaperone and Cooperates with ATRX in Replication-Independent Chromatin Assembly at Telomeres.” Proceedings of the National Academy of Sciences 107 (32) 14075–14080 https://doi.org/10.1073/pnas.1008850107.

Loppin, Benjamin, Emilie Bonnefoy, Caroline Anselme, Anne Laurençon, Timothy L. Karr, and Pierre Couble. 2005. “The Histone H3.3 Chaperone HIRA Is Essential for Chromatin Assembly in the Male Pronucleus.” Nature 437 (7063): 1386–90. https://doi.org/10.1038/nature04059.

Loyola, Bonaldi, Tiziana, Roche, Daniele, and Almouzni, Genevieve. 2006. “PTMs on H3 Variants BeforeChromatin Assembly PotentiateTheir Final Epigenetic State.” Molecular Cell 24(2):309–16. https://doi.org/10.1016/j.molcel.2006.08.019.

Lu, Huasong, Dan Yu, Anders S. Hansen, Sourav Ganguly, Rongdiao Liu, Alec Heckert, Xavier Darzacq, and Qiang Zhou. 2018. “Phase-Separation Mechanism for C-Terminal Hyperphosphorylation of RNA Polymerase II.” Nature 558 (7709): 318–23. https://doi.org/10.1038/s41586-018-0174-3.

Luger, Karolin. 1997. “Crystal Structure of the Nucleosome Core Particle at 2.8 Å Resolution” Proceedings of the National Academy of Sciences 107(32):14075–80. https://doi.org/10.1073/pnas.1008850107.

Martini, Emmanuelle, Danièle M.J. Roche, Kathrin Marheineke, Alain Verreault, and Geneviève Almouzni. 1998. “Recruitment of Phosphorylated Chromatin Assembly Factor 1 to Chromatin after UV Irradiation of Human Cells.” The Journal of Cell Biology 143 (3): 563–75. https://doi.org/10.1083/jcb.143.3.563.

Mayer, Andreas, Michael Lidschreiber, Matthias Siebert, Kristin Leike, Johannes Söding, and Patrick Cramer. 2010. “Uniform Transitions of the General RNA Polymerase II Transcription Complex.” Nature Structural & Molecular Biology 17 (10): 1272–78. https://doi.org/10.1038/nsmb.1903.

Maze, Ian, Wendy Wenderski, Kyung-Min Noh, Rosemary C. Bagot, Nikos Tzavaras, Immanuel Purushothaman, Simon J. Elsässer, et al. 2015. “Critical Role of Histone Turnover in Neuronal Transcription and Plasticity.” Neuron 87 (1): 77–94. https://doi.org/10.1016/j.neuron.2015.06.014.

Mello, J. A. 2002. “Human Asf1 and CAF-1 Interact and Synergize in a Repair-Coupled Nucleosome Assembly Pathway.” EMBO Reports 3 (4): 329–34. https://doi.org/10.1093/embo-reports/kvf068.

Nagashima, Ryosuke, Kayo Hibino, S.S. Ashwin, Michael Babokhov, Shin Fujishiro, Ryosuke Imai, Tadasu Nozaki, et al. 2019. “Single Nucleosome Imaging Reveals Loose Genome Chromatin Networks via Active RNA Polymerase II.” The Journal of Cell Biology 218 (5): 1511–30. https://doi.org/10.1083/jcb.201811090.

Nourani, A., F. Robert, and F. Winston. 2006. “Evidence That Spt2/Sin1, an HMG-Like Factor, Plays Roles in Transcription Elongation, Chromatin Structure, and Genome Stability in Saccharomyces Cerevisiae.” Molecular and Cellular Biology 26 (4): 1496–1509. https://doi.org/10.1128/MCB.26.4.1496-1509.2006.

Nozaki, Tadasu, Ryosuke Imai, Mai Tanbo, Ryosuke Nagashima, Sachiko Tamura, Tomomi Tani, Yasumasa Joti, et al. 2017. “Dynamic Organization of Chromatin Domains Revealed by Super-Resolution Live-Cell Imaging.” Molecular Cell 67 (2): 282–293.e7. https://doi.org/10.1016/j.molcel.2017.06.018.

Orsi, Guillermo A, Ahmed Algazeery, Meyer, Capri, and Sapey. 2013. “Drosophila Yemanuclein and HIRA Cooperate for De Novo Assembly of H3.3-Containing Nucleosomes in the Male Pronucleus.” PLOS Genetics 9 (2): 13.

Otterstrom, Jason, Alvaro Castells-Garcia, Chiara Vicario, Pablo A Gomez-Garcia, Maria Pia Cosma, and Melike Lakadamyali. 2019. “Super-Resolution Microscopy Reveals How Histone Tail Acetylation Affects DNA Compaction within Nucleosomes in Vivo.” Nucleic Acids Research, July, gkz593. https://doi.org/10.1093/nar/gkz593.

Pchelintsev, Nikolay A., Tony McBryan, Taranjit Singh Rai, John van Tuyn, Dominique Ray-Gallet, Geneviève Almouzni, and Peter D. Adams. 2013. “Placing the HIRA Histone Chaperone Complex in the Chromatin Landscape.” Cell Reports 3 (4): 1012–19. https://doi.org/10.1016/j.celrep.2013.03.026.

Petesch, Steven J., and John T. Lis. 2008. “Rapid, Transcription-Independent Loss of Nucleosomes over a Large Chromatin Domain at Hsp70 Loci.” Cell 134 (1): 74–84. https://doi.org/10.1016/j.cell.2008.05.029.

Quivy, J.-P. 2001. “Dimerization of the Largest Subunit of Chromatin Assembly Factor 1: Importance in Vitro and during Xenopus Early Development.” The EMBO Journal 20 (8): 2015–27. https://doi.org/10.1093/emboj/20.8.2015.

Rai, T. S., A. Puri, T. McBryan, J. Hoffman, Y. Tang, N. A. Pchelintsev, J. van Tuyn, R. Marmorstein, D. C. Schultz, and P. D. Adams. 2011. “Human CABIN1 Is a Functional Member of the Human HIRA/UBN1/ASF1a Histone H3.3 Chaperone Complex.” Molecular and Cellular Biology 31 (19): 4107–18. https://doi.org/10.1128/MCB.05546-11.

Ray-Gallet, Dominique, M. Daniel Ricketts, Yukari Sato, Kushol Gupta, Ekaterina Boyarchuk, Toshiya Senda, Ronen Marmorstein, and Geneviève Almouzni. 2018. “Functional Activity of the H3.3 Histone Chaperone Complex HIRA Requires Trimerization of the HIRA Subunit.” Nature Communications 9 (1): 3103. https://doi.org/10.1038/s41467-018-05581-y.

Ray-Gallet, Dominique, Adam Woolfe, Isabelle Vassias, Céline Pellentz, Nicolas Lacoste, Aastha Puri, David C. Schultz, et al. 2011. “Dynamics of Histone H3 Deposition In Vivo Reveal a Nucleosome Gap-Filling Mechanism for H3.3 to Maintain Chromatin Integrity.” Molecular Cell 44 (6): 928–41. https://doi.org/10.1016/j.molcel.2011.12.006.

Ricci, Maria Aurelia, Carlo Manzo, María Filomena García-Parajo, Melike Lakadamyali, and Maria Pia Cosma. 2015. “Chromatin Fibers Are Formed by Heterogeneous Groups of Nucleosomes In Vivo.” Cell 160 (6): 1145–58. https://doi.org/10.1016/j.cell.2015.01.054.

Ricketts, M Daniel, Nirmalya Dasgupta, Jiayi Fan, Joseph Han, Morgan Gerace, Yong Tang, Ben E Black, Peter D Adams, and Ronen Marmorstein. 2019. “The HIRA Histone Chaperone Complex Subunit UBN1 Harbors H3/H4- and DNA-Binding Activity.” Journal of Biological Chemistry 294(23):9239–9259. https://doi.org/10.1074/jbc.RA119.007480

Ricketts, M Daniel, Brian Frederick, Henry Hoff, Yong Tang, David C Schultz, Taranjit Singh Rai, Maria Grazia Vizioli, Peter D Adams, and Ronen Marmorstein. 2015. “Ubinuclein-1 Confers Histone H3.3-Specific-Binding by the HIRA Histone Chaperone Complex.” Nature communications, 10;6:7711. https://doi.org/10.1038/ncomms8711.

Sabari, Benjamin R., Alessandra Dall’Agnese, Ann Boija, Isaac A. Klein, Eliot L. Coffey, Krishna Shrinivas, Brian J. Abraham, et al. 2018. “Coactivator Condensation at Super-Enhancers Links Phase Separation and Gene Control.” Science 361 (6400): eaar3958. https://doi.org/10.1126/science.aar3958.

Schneiderman, J. I., G. A. Orsi, K. T. Hughes, B. Loppin, and K. Ahmad. 2012. “Nucleosome-Depleted Chromatin Gaps Recruit Assembly Factors for the H3.3 Histone Variant.” Proceedings of the National Academy of Sciences 109 (48): 19721–26. https://doi.org/10.1073/pnas.1206629109.

Schwabish, M. A., and K. Struhl. 2004. “Evidence for Eviction and Rapid Deposition of Histones upon Transcriptional Elongation by RNA Polymerase II.” Molecular and Cellular Biology 24 (23): 10111–17. https://doi.org/10.1128/MCB.24.23.10111-10117.2004.

Schwartz, B. E., and Ahmad. 2005. “Transcriptional Activation Triggers Deposition and Removal of the Histone Variant H3.3.” Genes & Development 19 (7): 804–14. https://doi.org/10.1101/gad.1259805.

Shaban, Haitham A, Roman Barth, and Kerstin Bystricky. 2018. “Formation of Correlated Chromatin Domains at Nanoscale Dynamic Resolution during Transcription.” Nucleic Acids Research 46 (13): e77–e77. https://doi.org/10.1093/nar/gky269.

Simon, Aline C., Jin C. Zhou, Rajika L. Perera, Frederick van Deursen, Cecile Evrin, Marina D. Ivanova, Mairi L. Kilkenny, et al. 2014. “A Ctf4 Trimer Couples the CMG Helicase to DNA Polymerase α in the Eukaryotic Replisome.” Nature 510 (7504): 293–97. https://doi.org/10.1038/nature13234.

Svensson, J. Peter, Manu Shukla, Victoria Menendez-Benito, Ulrika Norman-Axelsson, Pauline Audergon, Indranil Sinha, Jason C. Tanny, Robin C. Allshire, and Karl Ekwall. 2015. “A Nucleosome Turnover Map Reveals That the Stability of Histone H4 Lys20 Methylation Depends on Histone Recycling in Transcribed Chromatin.” Genome Research 25 (6): 872–83. https://doi.org/10.1101/gr.188870.114.

Tagami, Hideaki, Dominique Ray-Gallet, Geneviève Almouzni, and Yoshihiro Nakatani. 2004. “Histone H3.1 and H3.3 Complexes Mediate Nucleosome Assembly Pathways Dependent or Independent of DNA Synthesis.” Cell 116 (1): 51–61. https://doi.org/10.1016/S0092-8674(03)01064-X.

Talbert, Paul B., and Steven Henikoff. 2010. “Histone Variants — Ancient Wrap Artists of the Epigenome.” Nature Reviews Molecular Cell Biology 11 (4): 264–75. https://doi.org/10.1038/nrm2861.

Tang, Yong, Maxim V Poustovoitov, Kehao Zhao, Megan Garfinkel, Adrian Canutescu, Roland Dunbrack, Peter D Adams, and Ronen Marmorstein. 2006. “Structure of a Human ASF1a–HIRA Complex and Insights into Specificity of Histone Chaperone Complex Assembly.” Nature Structural & Molecular Biology 13 (10): 921–29. https://doi.org/10.1038/nsmb1147.

Teves, Sheila S, and Steven Henikoff. 2014. “Transcription-Generated Torsional Stress Destabilizes Nucleosomes.” Nature Structural & Molecular Biology 21 (1): 88–94. https://doi.org/10.1038/nsmb.2723.

Thebault, P., G. Boutin, W. Bhat, A. Rufiange, J. Martens, and A. Nourani. 2011. “Transcription Regulation by the Noncoding RNA SRG1 Requires Spt2-Dependent Chromatin Deposition in the Wake of RNA Polymerase II.” Molecular and Cellular Biology 31 (6): 1288–1300. https://doi.org/10.1128/MCB.01083-10.

Thiriet, C., and Jeffrey J. Hayes. 2005. “Replication-Independent Core Histone Dynamics at Transcriptionally Active Loci in Vivo.” Genes & Development 19 (6): 677–82. https://doi.org/10.1101/gad.1265205.

Torné, Júlia, Guillermo A. Orsi, Dominique Ray-Gallet, and Geneviève Almouzni. 2018. “Imaging Newly Synthesized and Old Histone Variant Dynamics Dependent on Chaperones Using the SNAP-Tag System.” In Histone Variants, edited by Guillermo A. Orsi and Geneviève Almouzni, 1832:207–21. New York, NY: Springer New York. https://doi.org/10.1007/978-1-4939-8663-7_11.

Tyler, Jessica K., Christopher R. Adams, Shaw-Ree Chen, Ryuji Kobayashi, Rohinton T. Kamakaka, and James T. Kadonaga. 1999. “The RCAF Complex Mediates Chromatin Assembly during DNA Replication and Repair.” Nature 402 (6761): 555–60. https://doi.org/10.1038/990147.

Venkatesh, Swaminathan, and Jerry L. Workman. 2015. “Histone Exchange, Chromatin Structure and the Regulation of Transcription.” Nature Reviews Molecular Cell Biology 16 (3): 178–89. https://doi.org/10.1038/nrm3941.

Vos, Seychelle M., Lucas Farnung, Marc Boehning, Christoph Wigge, Andreas Linden, Henning Urlaub, and Patrick Cramer. 2018. “Structure of Activated Transcription Complex Pol II–DSIF–PAF–SPT6.” Nature 560 (7720): 607–12. https://doi.org/10.1038/s41586-018-0440-4.

Wong, L. H., J. D. McGhie, M. Sim, M. A. Anderson, S. Ahn, R. D. Hannan, A. J. George, K. A. Morgan, J. R. Mann, and K.H. A. Choo. 2010. “ATRX Interacts with H3.3 in Maintaining Telomere Structural Integrity in Pluripotent Embryonic Stem Cells.” Genome Research 20 (3): 351–60. https://doi.org/10.1101/gr.101477.109.

Yadav, Tejas, Jean-Pierre Quivy, and Geneviève Almouzni. 2018. “Chromatin Plasticity: A Versatile Landscape That Underlies Cell Fate and Identity.” Science 361 (6409): 1332–36. https://doi.org/10.1126/science.aat8950.

Yahli Lorch, Janice W. LaPointe, and Roger D. Kornberg. 1987. “Nucleosomes Inhibit the Initiation of Transcription but Allow Chain Elongation with the Displacement of Histones.” Cell 49(2):203–10.

Yoh, S. M., J. S. Lucas, and K. A. Jones. 2008. “The Iws1:Spt6:CTD Complex Controls Cotranscriptional MRNA Biosynthesis and HYPB/Setd2-Mediated Histone H3K36 Methylation.” Genes & Development 22 (24): 3422–34. https://doi.org/10.1101/gad.1720008.

Zhang, Gan, Wang, and Wold. 2017. “RPA Interacts with HIRA and Regulates H3.3 Deposition at Gene Regulatory Elements in Mammalian Cells.” Molecular Cell 65(2):272–284. https://doi.org/10.1016/j.molcel.2016.11.030.

